# The BOD1L subunit of the SETD1A complex sustains the expression of DNA damage repair genes despite restraining H3K4 trimethylation

**DOI:** 10.1101/2023.04.06.535882

**Authors:** Giovanni Ciotta, Sukhdeep Singh, Andrea Kranz, Ashish Gupta, Davi Coe Torres, Jun Fu, Rupam Choudhury, Nico Liske, Wai Kit Chu, Chuna Choudhary, Lenka Gahurova, Dmitrii Severinov, Ali Al-Fatlawi, Michael Schroeder, Rein Aasland, Konstantinos Anastassiadis, Anna R. Poetsch, A. Francis Stewart

## Abstract

SETD1A is the histone 3 lysine 4 (H3K4) methyltransferase central to the mammalian version of the highly conserved eight subunit Set1 complex (Set1C) that apparently conveys H3K4 trimethylation (H3K4me3) onto all active Pol II promoters. Accordingly, the *Setd1a* mouse knock-out dies in early embryogenesis at the epiblast stage and mouse embryonic stem cells (ESCs) die when SETD1A is removed. We report that ESC death is accompanied by loss of expression of DNA repair genes and accumulating DNA damage. BOD1L and BOD1 are homologs of the yeast Set1C subunit, Shg1, and subunits of the mammalian SETD1A and B complexes. We show that the Shg1 homology region binds to a highly conserved central alpha-helix in SETD1A and B. Like mutagenesis of *Shg1* in yeast, conditional mutagenesis of *Bod1l* in ESCs promoted increased H3K4 di- and tri-methylation but also, like loss of SETD1A, loss of expression of DNA repair genes, increased DNA damage and cell death. In contrast to similar losses of DNA repair gene expression, the converse changes in H3K4 methylation after loss of SETD1A or BOD1L implies that H3K4 methylation is not essential for expression of the target DNA repair genes. Because BOD1L becomes highly phosphorylated after DNA damage and acts to protect damaged replication forks, the SETD1A complex, and BOD1L in particular, are key nodes for the DNA damage repair network.

## Introduction

Post-translational modifications on histone 3 (H3) are principal determinants of chromatin status. In particular, methylation of H3 lysine 4 (H3K4me) is the primary characteristic of euchromatin and H3K4 trimethylation (H3K4me3) is a ubiquitous modification on nucleosomes at active promoters (Piunti and Shilatifard 2016; Howe et al. 2017; Talbert and Henikoff 2021). The first H3K4 methyltransferase complex to be identified, Set1C, was isolated from yeast. It contains eight subunits and was also the first linkage between H3K4 methylation and Trithorax-Group (TrxG) action (Roguev et al. 2001; Krogan et al. 2002). Concordant with the ubiquity of H3K4 methylation for eukaryotic euchromatin, Set1C is amongst the most highly conserved complexes in epigenetics. Mammals have two Set1Cs, based on the paralogs, SETD1A/KMT2F and SETD1B/KMT2G, and both complexes include highly conserved homologs of the other seven yeast Set1C subunits (Lee et al. 2007; van Nuland et al. 2013; Kranz and Anastassiadis 2020). Methylation is catalyzed by the SET domain of the largest subunit, SET1, however enzymatic activity also requires three of the subunits (Roguev et al. 2001; Morillon et al. 2005; Schneider et al. 2005; Dehe et al. 2006; Kim et al. 2013b). These three subunits (WDR5, RBBP5 and ASH2L) together with DPY30, form the highly conserved WRAD scaffold that is also common to the four mammalian MLL (MLL1-4/KMT2A-D) SET domain H3K4 methyltransferase complexes (Ernst and Vakoc 2012).

Although they share the same eight subunit architecture as yeast Set1C and are both probably dimeric (Choudhury et al. 2019), the mammalian SETD1A and SETD1B complexes (SETD1A-C, SETD1B-C) are distinct. No heteromeric complex containing both SETD1A and B has been detected (Lee and Skalnik 2005; Lee et al. 2007; Lee and Skalnik 2008; van Nuland et al. 2013). This concords with their distinct functions as revealed by knockout mice. *Setd1a* is required for the embryo to progress beyond the epiblast, whereas *Setd1b* knockout embryos proceed past gastrulation and die during organogenesis (Bledau et al. 2014). Furthermore, loss of SETD1A in mouse embryonic stem cells (ESCs) led to cell death whereas loss of SETD1B was tolerated and overexpression of SETD1B did not rescue the loss of SETD1A (Bledau et al. 2014).

Because the same six subunits (the four WRAD subunits plus CXXC1/CFP1 and WDR82) are components of both complexes, the functional differences between SETD1A-C and SETD1B-C must either be due to differences between SETD1A and SETD1B or differences between the eighth subunit. In yeast, the eighth subunit is Shg1 (Roguev et al. 2001; Roguev et al. 2003). The Shg1 subunit was originally missed in yeast COMPASS (Miller et al. 2001) and in the first mammalian and *Drosophila* SET1C/COMPASS descriptions (Lee and Skalnik 2005; Lee et al. 2007; Lee and Skalnik 2008; Ardehali et al. 2011; Mohan et al. 2011; Weiner et al. 2012). However, a mass spectrometry screen in HeLa cells identified two mammalian Shg1 homologues, BOD1 and BOD1L, as candidate SET1C subunits. BOD1 was identified as a candidate SETD1B-C subunit and BOD1L was identified as a candidate SETD1A-C subunit (van Nuland et al. 2013).

In yeast, Shg1 is not required for Set1C assembly and integrity but notably is the only Set1C subunit that restrains H3K4 trimethylation both *in vivo* (Roguev et al. 2001; Shevchenko et al. 2002; Roguev et al. 2003; Dehe et al. 2006) and *in vitro* (Kim et al. 2013b). That is, in the absence of Shg1, the remaining seven subunits of Set1C deliver elevated di- and tri-H3K4 methylation. BOD1 (<u>bio</u>rientation <u>d</u>eficient 1) was first found as a novel kinetochore protein required during mitotic chromosome segregation through action as an inhibitor of Protein Phosphatase 2A (PP2A) (Porter et al. 2007; Porter et al. 2013). BOD1L was first identified as a target of phosphorylation by ATM/ATR kinases after the induction of DNA damage (Matsuoka et al. 2007; Beli et al. 2012). Subsequently, a role in the protection of damaged replication forks was identified (Higgs et al. 2015). Shg1, BOD1L and BOD1 share the 110 a.a. Shg1 homology region that AlphaFold predicts as a bi-layered α-helical coat hanger (Fig. 1), suggesting that it serves as a binding adaptor, which is consistent with the role of BOD1 as an inhibitor of PP2A holoenzyme through subunit B56 (Schleicher et al. 2017).

**Figure 1.**
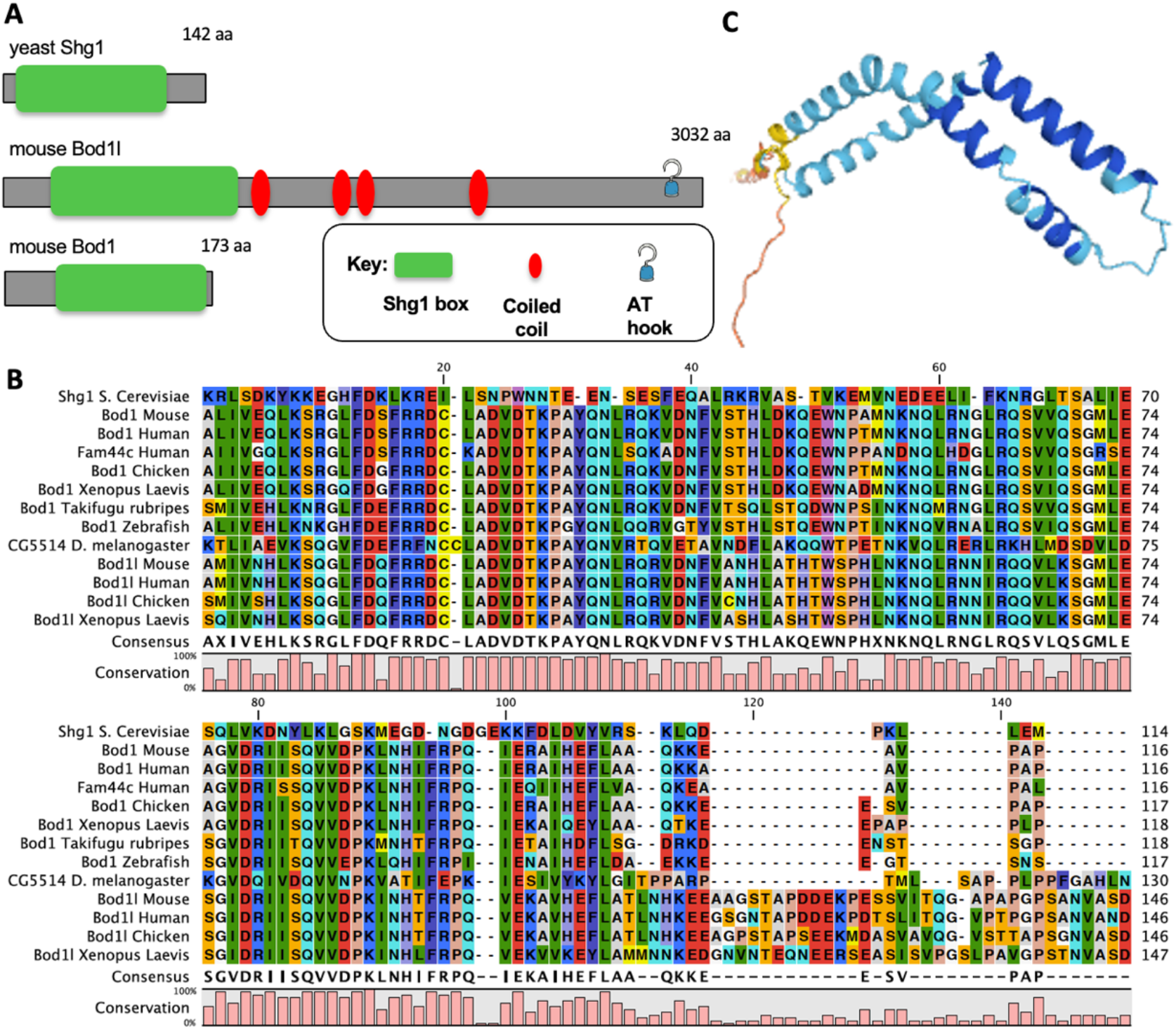
The Shg1 box in mammals. **A.** Protein diagrams of yeast Shg1, BOD1L and BOD1. **B.** Sequence alignment of the Shg1 box. Note humans have a third gene with a Shg1 box, originally designated Fam44c and now called BOD1L2 although it is, like BOD1 and unlike BOD1L, only slightly longer than the Shg1 box at 172 a.a.s. Also note that the BOD1L Shg1 box has an additional conserved C-terminal extension. **C.** AlphaFold prediction of the BOD1L Shg1 box. Dark blue – very high confidence; light blue – high confidence; yellow low confidence.

We uncovered the role of BOD1L in SETD1A-C in pursuit of the question: Why do mouse ESCs die shortly after conditional removal of SETD1A? We found that SETD1A-C and specifically BOD1L are required to maintain expression of genes involved in DNA repair. In their absence, accumulating DNA damage leads to induction of the DNA damage response and death. A similar relationship between SETD1A-C and DNA damage repair gene expression emerged from investigations of SETD1A-C in hematopoiesis (Arndt et al. 2018) and a leukemia cell line in culture (Hoshii et al. 2018). Here we identify BOD1L as the key SETD1A-C subunit in the central relationship between SETD1A-C and DNA damage repair gene expression.

## Results

### Characterizing the SET1 complexes in ESCs

To understand why ESCs die shortly after removal of SETD1A, we characterized the ESC SETD1A complex (SETD1A-C) using a *Setd1a* BAC transgene to express Venus-tagged SETD1A and a standardized GFP affinity purification and mass spectrometry protocol (AP-MS; (Choudhary and Mann 2010; Hubner et al. 2010; Hofemeister et al. 2011) to identify candidate interacting proteins. SETD1A AP-MS yielded most of the expected SET1C including RBBP5, ASH2L, DPY30, CXXC1, WDR82 and also BOD1L. HCFC1, which has been previously shown to associate with human SETD1A (Wysocka et al. 2003; Lee and Skalnik 2008) was also identified (Fig. 2A).

**Figure 2.**
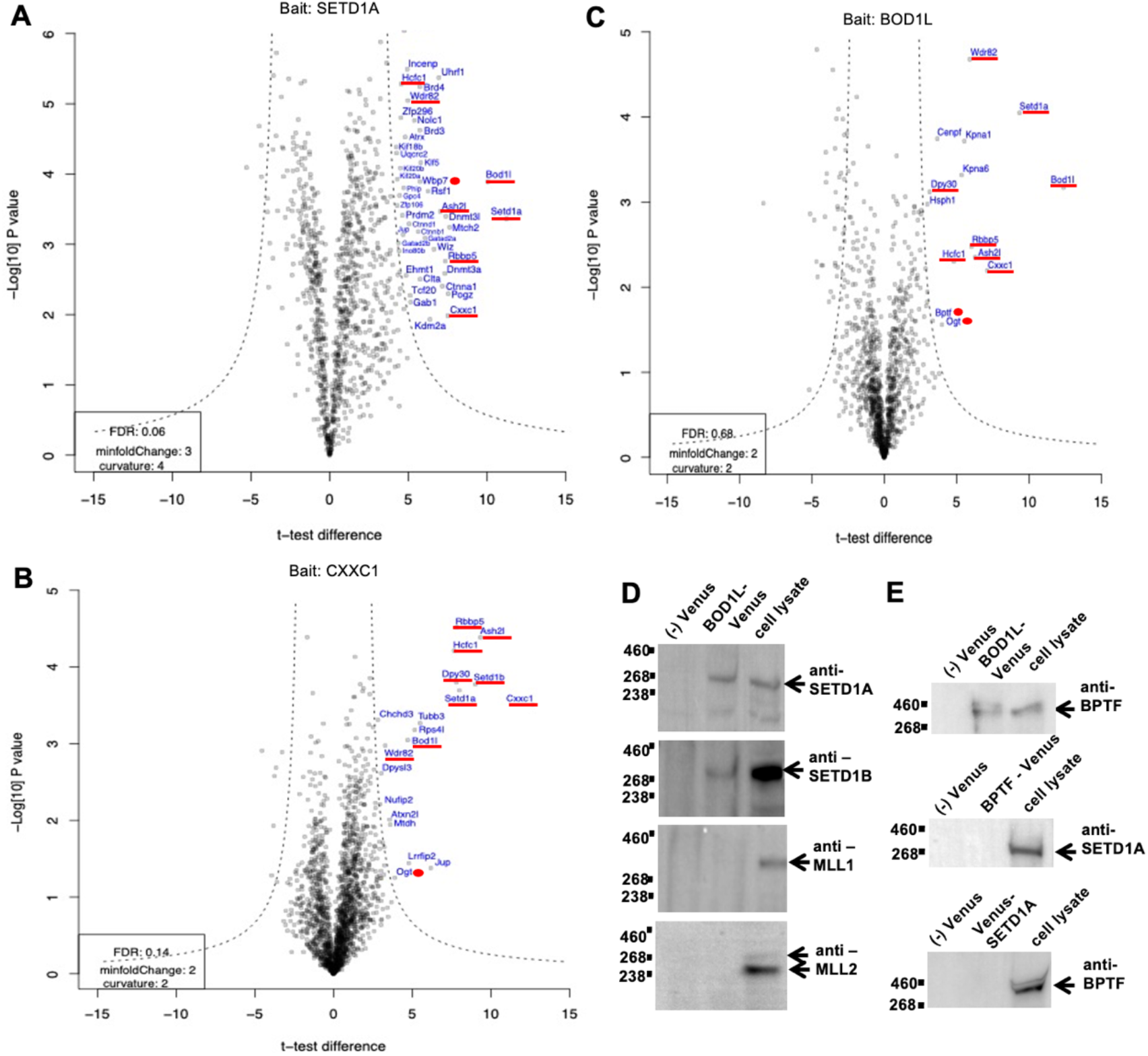
BOD1L is a subunit of the SETD1A complex in ESCs and also interacts with BPTF. **A.** Volcano plot of Venus-SETD1A AP-MS. Red underline denotes expected SETD1A-C subunits. Red dots indicate notable additional proteins (Wbp7 = Mll2/Kmt2b). **B.** As for A except CXXC1-Venus. **C.** As for A except BOD1L-Venus. **D.** BOD1L-Venus immunoprecipitation and Western analysis using anti-SETD1A, -SETD1B, -MLL1 and MLL2 antibodies. Lane (-) Venus, mock immunoprecipitation without anti-GFP antibody; lane BOD1L-Venus, immunoprecipitation with anti-GFP antibody; lane cell lysate, 10% total cell lysate used for the immunoprecipitation. **E.** As for D except immunoprecipitation using BOD1L-Venus, BPTF-Venus or Venus-SETD1A BAC ESC lines and anti-BPTF and anti-SETD1A antibodies as indicated.

We next employed BAC transgenes to express Venus-tagged CXXC1/CFP1, which is a subunit common to both SETD1A-C and SETD1B-C (Lee et al. 2007). As expected, CXXC1-Venus AP-MS yielded both SETD1A and SETD1B, plus five of the expected SET1C as well as BOD1L and again HCFC1 (Fig. 2B).

Consequently, we tagged BOD1L with Venus and AP-MS retrieved SETD1A as well as five expected SETD1A-C subunits and HCFC1 (Fig. 2C). Immunoprecipitation using Venus-tagged BOD1L confirmed the association with SETD1A (Fig. 2D) but no BOD1L interaction was observed with MLL1, MLL2 (Fig. 2D) or MLL4 (Supplemental Fig. S1). However, a modest interaction with SETD1B was observed (Fig. 2D).

These AP-MS analyses identified a variety of other candidate interactors. Of the nuclear proteins that we examined further, either by tagging and AP-MS or immunoprecipitation, only OGT and BPTF yielded constructive data. AP-MS with Venus tagged OGT retrieved SETD1A but also many other likely interacting proteins so we did not pursue this line of enquiry. Whether the multi-functional OGT (Yang and Qian 2017), which in *Drosophila* has been identified as the Polycomb-Group (PcG) protein *super sex comb* (*sxc*; (Gambetta et al. 2009) is a PcG member because it interacts with SETD1A-C requires further evaluation. BPTF is part of the promoter-associated NURF complex (Tsukiyama and Wu 1995), contains a PHD finger that binds H3K4me3 (Li et al. 2006) and acts downstream of WDR5 (Wysocka et al. 2006). Set1Cs also contain a PHD finger (in Spp1/CXXC1) that binds H3K4me3 (Shi et al. 2007; Eberl et al. 2013). Given this concordance, we examined the interaction with BPTF in more detail. By immunoprecipitation using BOD1L-VENUS expressed from a *Bod1l* BAC transgene, the interaction between BOD1L and BPTF was confirmed. Immunoprecipitation using BPTF-VENUS expressed from a *Bptf* BAC transgene retrieved the NURF subunit SNF2L/SMARCA1 as expected (Supplemental Fig. S1) but failed to retrieve SETD1A (Fig. 2E) indicating that BOD1L independently interacts with both SETD1A-C and BPTF, and that BPTF interacts with BOD1L independently of its interaction with NURF.

To investigate BOD1L and BOD1 interactions further, we performed AP-MS with Venus-tagged SETD1B and BOD1 (Fig. 3). SETD1B AP-MS retrieved most of the expected complex. However, in concordance with the immunoprecipitation result (Fig. 2D), BOD1L rather than BOD1 was retrieved. We attribute the absence of BOD1 in this AP-MS to the technical difficulty involved in mass spectroscopic identification of this small protein, which possibly explains the previous failures to identify Shg1 homologs in yeast, fly and mammalian SET1Cs. In agreement with this explanation, AP-MS with tagged BOD1 retrieved SETD1B and four expected SETD1-C subunits, confirming the interaction between SETD1B and BOD1. Notably, neither SETD1B nor BOD1 AP-MS analyses identified HCFC1, suggesting that HCFC1 exclusively interacts with SETD1A-C. Immunoprecipitation using tagged BOD1 confirmed interactions with both SETD1A and SETD1B but not MLL1, MLL2 or BPTF (Fig. 3C). Amongst candidate BOD1 interactors identified by AP-MS, SGO2 (Shugoshin 2) is notable because BOD1 is required for localization of its paralog, SGO1, at kinetochores (Porter et al. 2013).

**Figure 3.**
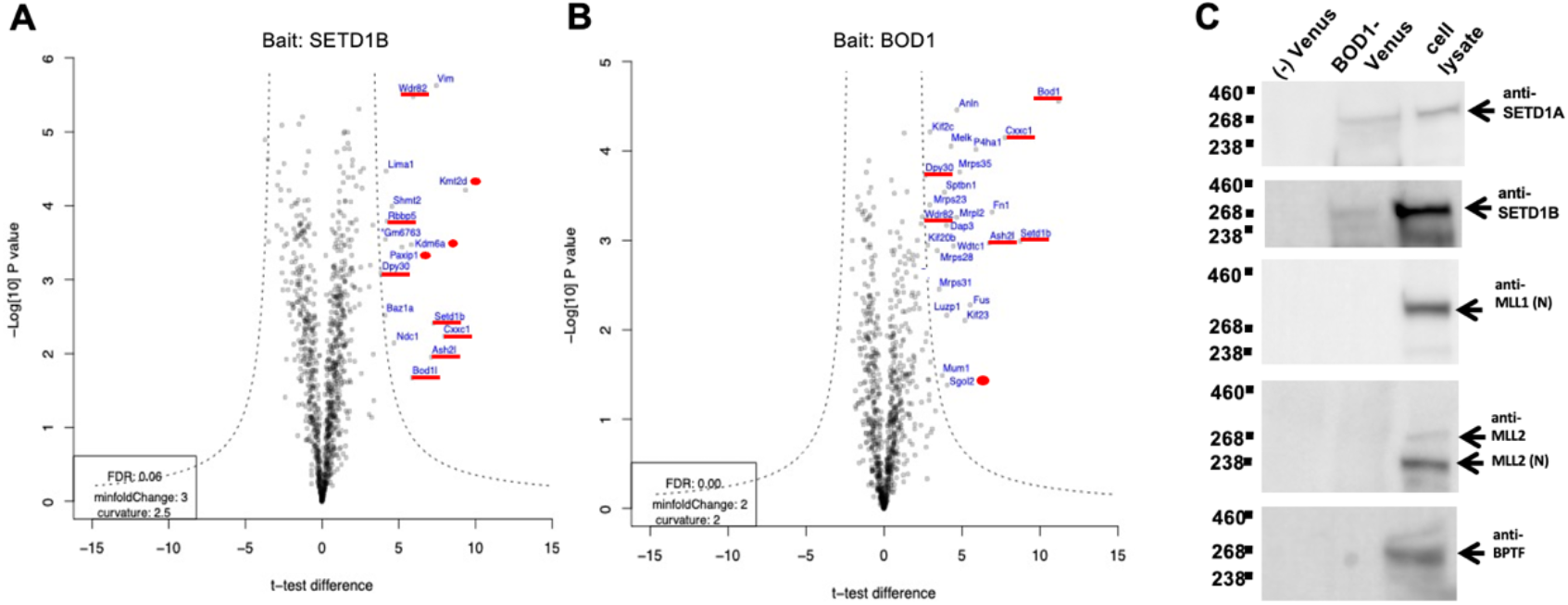
Both BOD1 and BOD1L associate with SETD1B and SETD1A in ESCs. **A.** As for Figure 2A except AP-MS with Venus-SETD1B. **B.** As for Figure 2A except AP-MS with BOD1-Venus. **C.** BOD1-Venus immunoprecipitation and Western analysis using anti-SETD1A, -SETD1B, -MLL1, -MLL2 and -BPTF antibodies. Lane (-) Venus, immunoprecipitation without anti-GFP antibody; lane BOD1-Venus, immunoprecipitation with anti-GFP antibody; lane cell lysate, 10% total cell lysate used for the immunoprecipitation.

### The Shg1 box interacts with a highly conserved helix in SET1

Because the Shg1 box is a highly conserved component of SET1Cs, we reasoned that its interaction site on SET1 will be highly conserved. SET1 sequence alignments across eukaryotic evolution originally defined the three most highly conserved domains; RRM, N-SET and SET plus the post-SET extension (Roguev et al. 2001). We identified further patches of homology in SET1 operationally termed X1 to X4 (Supplemental Fig. S2). Using the Venus-tagged *Setd1a* BAC transgene, we made versions deleted for X1, X3 and X4 (Supplemental Fig. S2) and analyzed associated proteins in ESCs by AP-MS (Fig. 4A-C). All three deletions retrieved the core SET1C subunits, RBBP5, DPY30, CXXC1 as well as HCFC1. Notably, BOD1L was retrieved in the X1 and X4 but not X3 deleted versions. Also, WDR82 was retrieved in the X3 and X4 but not X1 deleted versions, which is concordant with the WDR82 interaction site (Hughes et al. 2023).

**Figure 4.**
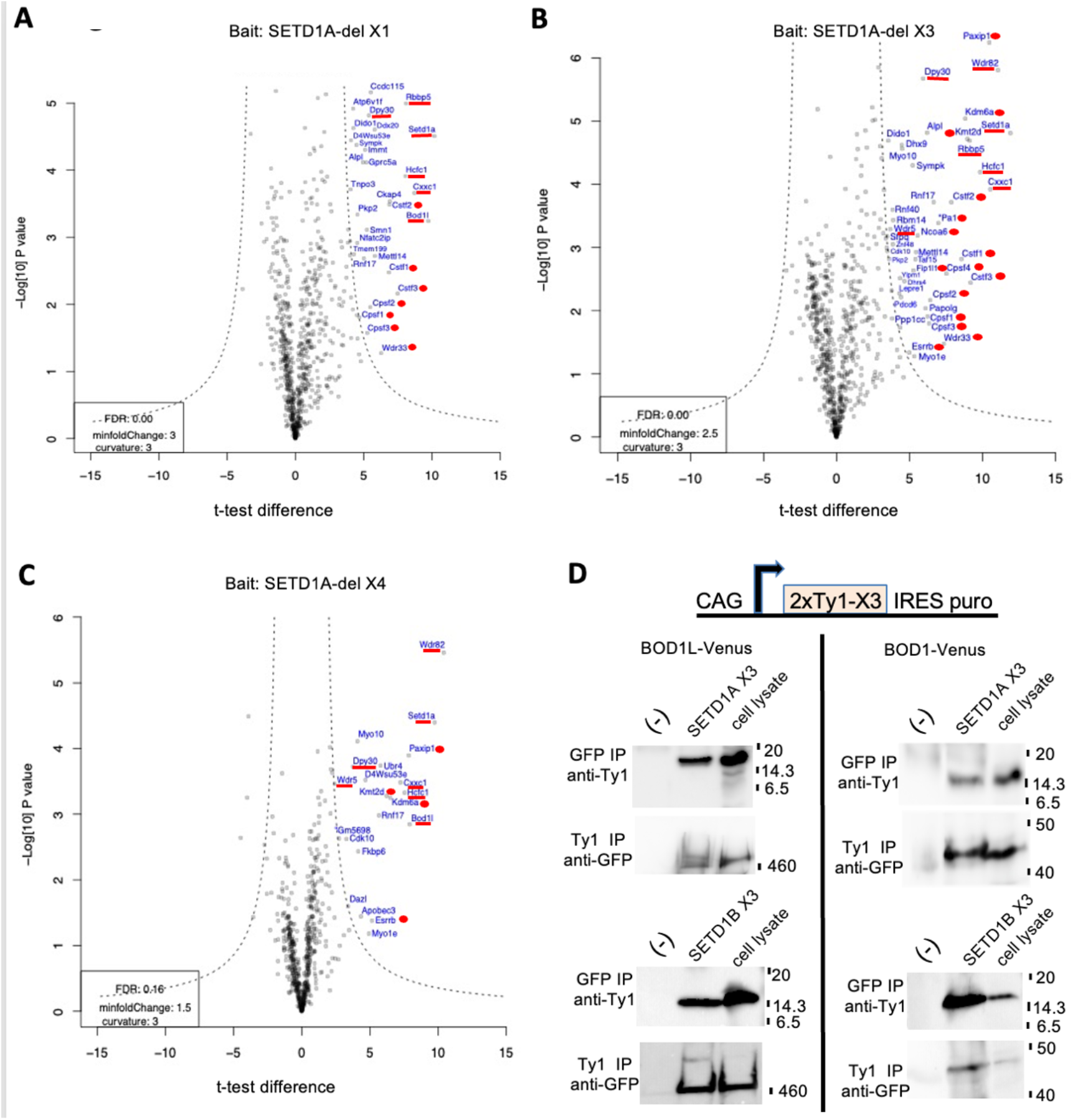
Both BOD1 and BOD1L interact with the X3 regions of SETD1A and SETD1B. **A-C.** As for Figure 2A except AP-MS with Venus-SETD1A with (**A**) X1, (**B**) X3, or (**C**) X4 region deleted. **D.** Immunoprecipitations from the ESCs expressing BOD1-Venus (left) or BOD1L-Venus were modified by introducing transgenes, selected for puromycin, expressing either 2xTy1-SETD1A-X3 (SETD1A a.a.s 787-874) or 2xTy1-SETD1B-X3 (SETD1B a.a.s 842-923) as illustrated. Immunoprecipitations were performed with either anti-GFP or anti-Ty1 antibodies and Westerns probed with the converse antibody. Lane (-) immunoprecipitation without anti-GFP or anti-Ty1 antibodies; lane SETD1A X3 (above) or SETD1B X3 (below) as indicated; lane cell lysate, 10% total cell lysate used for the immunoprecipitation.

We focused on the absence of BOD1L from the X3-deleted AP-MS analysis. X3 is a highly conserved SET1 region that is strongly predicted by AlphaFold to be a long α-helix (Supplemental Fig. S2). To test the interaction between BOD1L and the X3 regions at physiological expression levels, we stably expressed Ty1-tagged SETD1A-X3 and Ty1-tagged SETD1B-X3 in the ESCs expressing either BOD1L-Venus or BOD1-Venus. Immunoprecipitation with either GFP or Ty1 antibodies revealed that both X3 regions interact with both BOD1L and BOD1, which is consistent with the AP-MS and immunoprecipitation results presented in Figures 2 and 3. AlphaFold modelling of these interactions indicated an intimate fit between the two layered α-helical coat hanger and the SET1 X3 α-helices (Supplemental Fig. S3).

### BOD1L restrains H3K4 methylation and is required for ESC survival

To evaluate BOD1L function, we sequentially targeted both alleles to create *loxP* conditional ESCs, and then used ligand-regulated site specific recombination (Logie and Stewart 1995) to induce *Bod1l* knock-out upon administration of 4-OH tamoxifen (Schwenk et al. 1998) using CreERT2 (Feil et al. 1997) knocked into the *Rosa26* locus (Seibler et al. 2003) (Fig. 5A, B).

**Figure 5.**
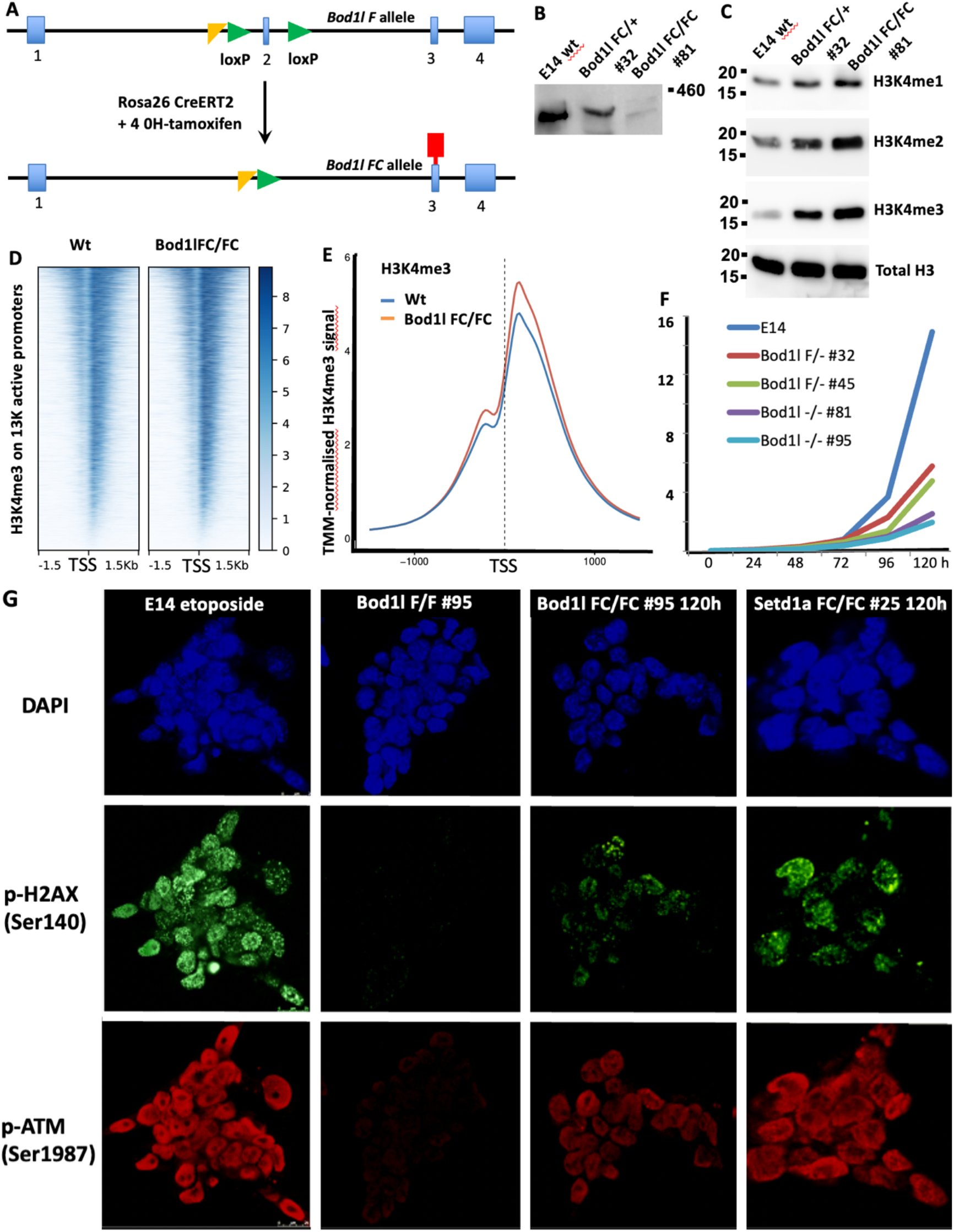
Conditional knock-out of *Bod1l* in ESCs. **A.** Diagram of the *Bod1l* loxP conditional allele (loxP sites denoted by green arrow heads). Removal of exon 2 causes a frame shift and stop codon in exon 3 (red block). Targeting was selected using an FRT flanked lacZ-neo cassette, which was removed by FLP recombination leaving an FRT site (yellow triangle). *CreERT2* was targeted into the *Rosa26* locus. **B.** Western blot probed with anti-BOD1L antibody evaluating - lane 1, E14 ESCs; lane 2, singly targeted, FLP and Cre recombined clone 32; lane 3, doubly targeted, FLP and Cre recombined clone 81. **C.** As for B except the Western was probed with antibodies against histone 3 and histone 3 lysine 4 methylations as indicated. **D.** Density plot of H3K4me3 ChIP-seq centered on transcription start sites (TSS) 3kb upstream and downstream for the 12.2 thousand active promoters in ESCs, ordered according to signal strength. **E.** As for E except accumulated signal strength. **F.** ESC proliferation after seeding 10^5^ cells of the indicated genotype. **G.** ESCs evaluated for DNA damage by staining for DAPI and phosphorylated H2A and ATM antibodies. Top row, E14 cells treated with etoposide (0.2μM, 24 hours); lower rows, *Bod1l* and *Setd1a* conditional ESCs treated with 4-hydroxy tamoxifen for 48 hours (indicated by FC) or not (F/F) then analyzed 72 hours later.

In both *S. cerevisiae* and *S. pombe*, Shg1 was the only Set1C subunit whose deletion resulted in increased global levels of H3K4me2 and 3 (Roguev et al. 2001; Roguev et al. 2003; Dehe et al. 2006). Remarkably, deletion of *Bod1l* in ESCs also provoked increased H3K4me2 and 3 (Fig. 5C, Supplemental Fig. S5B). H3K4me2 and 3 levels were not only elevated in *Bod1l-/-* ESCs but also increased about half as much in *Bod1l+/-* ESCs, indicating a dosage relationship between H3K4me2/3 and BOD1L protein expression levels. H3K4me2 and 3 levels were also increased in *Bod1l* esiRNAi knock-down experiments that reduced BOD1L to ∼25% (Supplemental Fig. S5A,B). Using ChIP-seq, we observed that after removal of BOD1L the bulk of the increased H3K4me3 occured on existing sites of H3K4me3, namely nucleosomes surrounding active promoters (Fig. 5D,E), indicating that BOD1L simply restricts methylation by SETD1A-C rather than influencing site selection. Taken together, these mass spectrometric, co-immunoprecipitation and functional data prove that BOD1L is a SETD1A-C subunit in ESCs and indicate that BOD1L expression levels closely restrict H3K4 di- and tri-methylation by SETD1A-C.

Notably, ESC doubling times were also prolonged in homozygous and heterozygous knock-outs according to gene dosage (Fig. 5F), which was not due to impact on the cell cycle (Supplemental Fig. S5C). Because of the relationships between BOD1L and DNA damage (Matsuoka et al. 2007; Beli et al. 2012; Higgs et al. 2015), we examined whether the loss of BOD1L compromised DNA damage repair. Unrepaired DNA damage accumulated shortly after loss of not only BOD1L but also SETD1A (Fig. 5G), which presents a straightforward explanation for the delayed growth and death of ESCs in both cases.

### BOD1L is the SETD1A-C subunit required for expression of many DNA repair genes

To evaluate the impact of the loss of BOD1L and SETD1A on gene expression, mRNA was harvested 96 hours after inducing *loxP* recombination with 4-OH tamoxifen. We chose this time point based on previous applications of *RosaCreERT2*/tamoxifen for conditional mutagenesis in ESCs (Bledau et al. 2014; Denissov et al. 2014), which revealed almost complete loss of target protein 48 hours after tamoxifen induction was initiated (corresponding to three to four cell cycles). Also, cell death of both *Setd1a* and *Bod1l* ESCs began four to five days after tamoxifen induction was initiated. For comparison, in parallel we evaluated tamoxifen-induced conditional mutagenesis of *Mll2/Kmt2b*, which does not lead to cell death (Denissov et al. 2014).

Three cellular conditions were evaluated in duplicates;

i. naïve ESCs cultured with LIF and 2i (two inhibitors; the MEK inhibitor, PD0325901: the GSK3β inhibitor, CT99021; (Ying et al. 2008; Marks et al. 2012);
ii. cycling ESCs cultured with FCS and LIF (Marks et al. 2012);
iii. ESCs differentiated towards epiblast-like stem cells (EpiSCs) by culture without LIF in the presence of Activin A and FGF2 (Hayashi et al. 2011).

Pairwise clustering of the RNA-seq data indicated high data quality (Supplemental Fig. S6A,B). Principle component analysis revealed that all duplicates clustered together and cellular state was more important than the absence of the target protein (Supplemental Fig. S6B). All three cellular states clustered separately with one exception. Both the uninduced and induced *Setd1a* EpiSC transcriptomes remained closer to the cycling ESC transcriptomes than to the EpiSC *Bod1l* and *Mll2/Kmt2b* transcriptomes, which is a result we attribute to incomplete transition of the *Setd1a* cells from cycling ESCs to EpiSCs, despite being cultured with Activin A and FGF2 in parallel with the *Bod1l* and *Mll2/Kmt2b* ESCs for this experiment. Because all three culture conditions delivered similar primary results (Supplemental Fig. S6C), this incomplete cellular transition to EpiSCs did not alter our conclusions.

In all three cell states, loss of either SETD1A or BOD1L resulted in substantially more reduced differentially expressed genes (DEGs) than increased DEGs, as illustrated in Volcano plots (Fig. 6A) and scatterplot analyses (Fig. 6B). This pronounced bias towards loss of gene expression concords with the expectation that SETD1A-C sustains transcriptional activity, and indicates that the observed DEGs are largely primary targets with few secondary perturbations.

**Figure 6.**
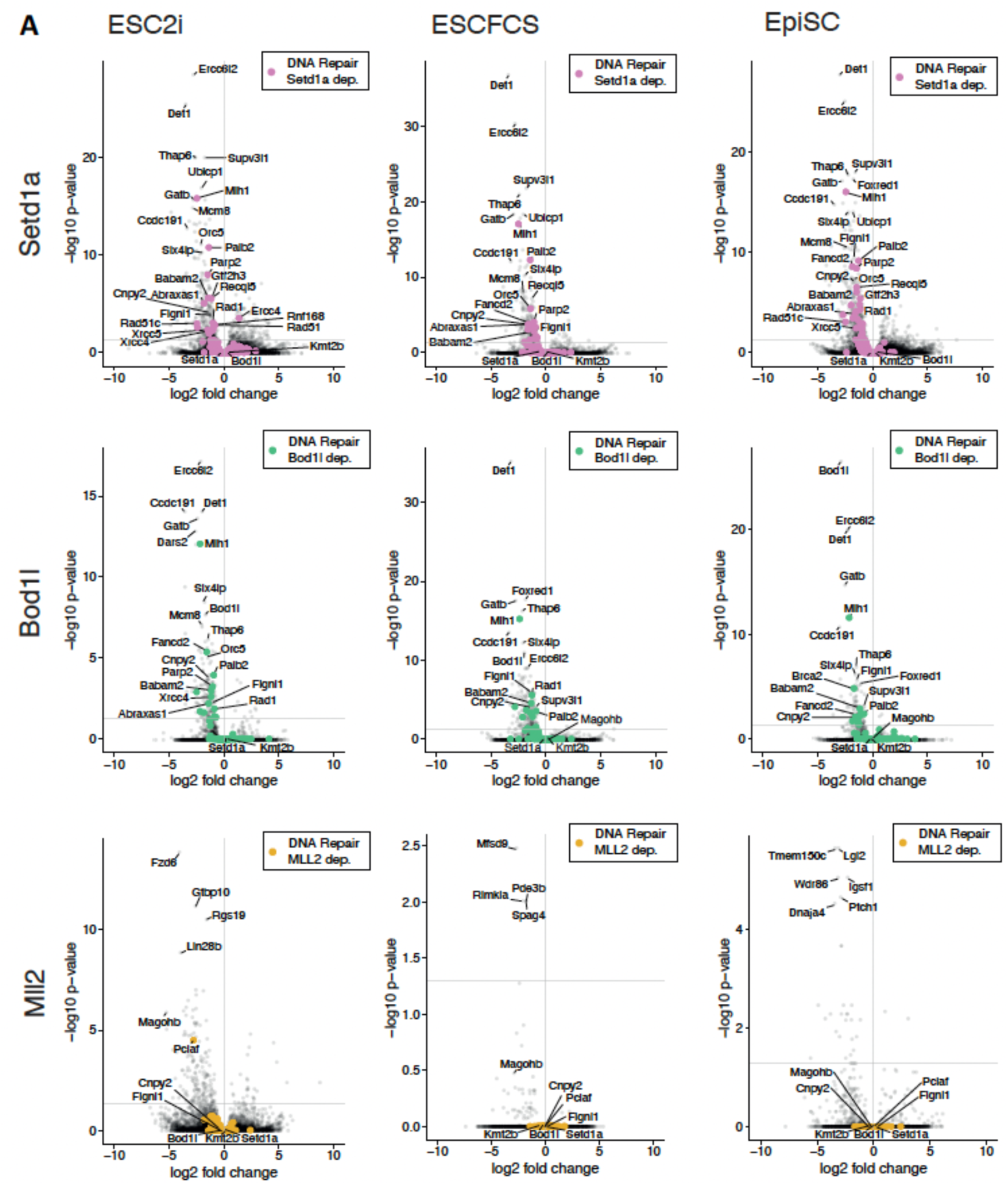
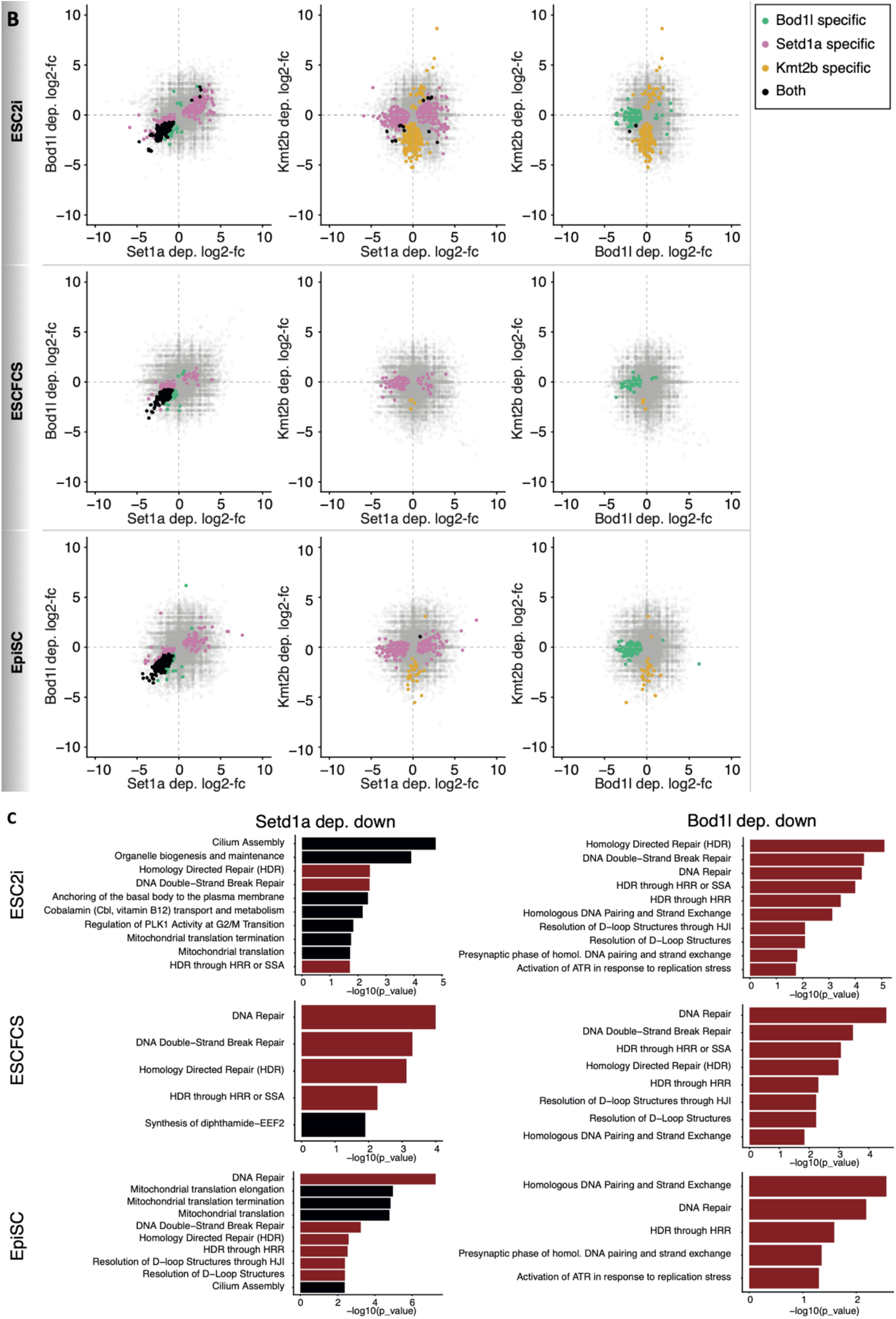
Transcriptome analysis of *Bod1l*, *Setd1a* and *Mll2* conditional knockouts. **A.** Volcano plots of the 36 RNA-seq datasets from the three cell types (ESC2i, ESCFCS, EpiSC) and the three genes *Setd1a, Bod1l, Mll2/Kmt2b* representing pairwise differential gene expression analyses dependent on 4-OH tamoxifen. DNA repair genes assigned by Reactome are denoted by coloured dots. *Ercc6l1* and *Slx4ip* were identified as DNA repair genes (Zhang et al. 2019; Olivieri et al. 2020) after the Reactome assignments used here so were not denoted with coloured dots **B.** Pairwise comparisons of gene expression from the three genes, *Setd1a, Bod1l, Mll2/Kmt2b* using conditional knock-out dependent log2 fold changes from the differential analysis. Genes are highlighted when significant with an adjusted p-value < 0.05 within the stated condition. For visibility, axes were capped at the absolute maximum value of the genes significant in any samples or at a log2-fold change of 10. **C.** GO term analysis with Reactome pathways of genes determined as differentially down regulated from the 36 datasets. Significant pathways with padj<0.05 are selected and highlighted in red when linked to ‘DNA Repair’. HRR – homologous recombination repair; SSA – single strand annealing; HJI – Holliday junction intermediates.

In all three culture conditions, the correlations of the effect sizes of differential expression revealed that *Bod1l* DEGs are subsets of *Setd1a* DEGs and these subsets are almost exclusively downregulated (Fig. 6B; Supplemental Fig S6C). In contrast, there was no correlation between *Setd1a* or *Bod1l* DEGs with *Mll2/Kmt2b* DEGs (Supplemental Fig. 6D). Consequently, we focused our analysis on *Setd1a* and *Bod1l* DEGs with decreased expression.

Decreased DEGs after loss of *Setd1a* presented a spectrum of functions with high concurrence between the three cell states. The most recurrent, significantly enriched, GO terms encompassed “DNA repair” and related subterms. Mitochondrial and cilium assembly terms, and Vitamin B12 metabolism were also prominent (Fig. 6C). Decreased expression of DNA repair genes included primarily components of homology directed repair (Table 1) indicating that SETD1A sustains this route for double strand break repair.

**Table 1.** DNA repair genes according to Reactome.db_ (version 1.82.0) with decreased expression (padj≤0.05) in at least two of the six cell states are listed with log2FoldChange. *Ercc6l1* and *Slx4ip* are DNA repair genes not included in Reactome.db_(version 1.82.0) and were added here.

|  | Setd1a | Setd1a | Setd1a | Bod1l | Bod1l | Bod1l |
| --- | --- | --- | --- | --- | --- | --- |
| Gene | ESC2i | ESCFCS | EpiSC | ESC2i | ESCFCS | EpiSC |
| Abraxas1 | -1,83 | -1,63 | -1,64 | -1,39 | -1,30 | -1,19 |
| Babam2 | -1,43 | -1,10 | -1,48 | -1,16 | -1,17 | -1,34 |
| Blm | -0,53 |  |  |  | -0,49 |  |
| Brca2 |  | -0,91 | -1,52 | -0,59 | -1,72 | -1,46 |
| Brip1 |  |  | -0,76 |  | -0,74 |  |
| Cenpx | -0,94 |  | -1,11 |  |  | -0,91 |
| Cetn2 | -0,91 |  | -0,93 |  |  |  |
| Cops3 | -0,66 | -0,89 | -1,13 |  |  |  |
| Dclre1b |  | -0,62 | -0,69 |  |  |  |
| Ddb2 |  |  | -0,54 | -0,54 |  |  |
| Eme1 | -0,76 | -1,12 | -1,35 | -0,69 |  | -0,58 |
| Ercc6l2 | -2,74 | -2,58 | -2,76 | -2,14 | -1,62 | -2,28 |
| Fan1 | -0,78 | -1,64 | -1,97 |  | -1,36 | -1,79 |
| Fancd2 | -0,69 | -1,33 | -1,85 | -1,58 | -1,16 | -1,22 |
| Fanci |  | -0,61 | -0,99 |  | -0,48 | -0,91 |
| Gen1 | -0,99 | -1,40 | -1,48 | -0,87 | -1,05 | -0,73 |
| Gtf2h3 | -1,16 | -0,95 | -1,14 | -0,73 | -0,46 | -0,67 |
| Gtf2h4 |  |  | -1,04 |  |  | -1,26 |
| Mcrs1 | -0,56 |  | -0,96 |  |  |  |
| Mlh1 | -2,47 | -2,51 | -2,45 | -2,18 | -2,15 | -2,40 |
| Mre11a | -0,44 | -0,41 |  |  | -0,45 |  |
| Neil1 | -1,06 |  | -0,84 | -0,86 |  | -0,79 |
| Neil2 | -1,94 |  |  | -1,34 |  |  |
| Nhej1 |  | -1,05 | -1,12 |  |  | -0,79 |
| Palb2 | -1,38 | -1,45 | -1,29 | -0,93 | -0,79 | -0,90 |
| Parp2 | -1,48 | -1,10 | -1,49 | -1,04 | -0,67 | -0,64 |
| Pold1 | -0,65 |  | -0,82 |  |  |  |
| Pold2 | -0,79 |  | -1,05 |  |  |  |
| Pold4 |  |  | -1,13 |  |  | -1,21 |
| Poli | -1,18 | -1,12 | -1,31 | -1,25 | -1,41 | -1,68 |
| Poll | -1,15 | -1,44 | -1,82 | -0,88 |  | -1,21 |
| Prkdc | -1,08 |  | -0,75 | -1,24 | -1,58 | -0,82 |
| Rad1 | -1,00 | -0,84 | -1,18 | -0,87 | -0,44 | -1,28 |
| Rad17 | -0,52 | -0,69 | -0,51 |  |  |  |
| Rad18 |  | -0,67 | -0,99 |  |  |  |
| Rad23a | -1,35 | -0,88 | -1,11 |  |  |  |
| Rad51 | -0,83 | -0,43 | -0,57 |  |  |  |
| Rad51b | -2,41 | -1,72 | -2,72 | -2,49 | -1,68 | -2,83 |
| Rad51c | -2,46 | -1,95 | -2,49 | -2,19 | -1,42 | -1,90 |
| Rad9b | -1,16 | -1,46 | -1,53 | -1,82 | -1,86 | -2,12 |
| Rbx1 |  | -0,62 | -1,08 |  |  |  |
| Rfc4 | -0,93 |  | -0,78 | -0,59 |  |  |
| Rfc5 |  |  | -0,91 |  |  | -0,49 |
| Rnf168 | -0,89 | -0,80 | -0,51 | -0,42 | -0,47 |  |
| Slx4ip | -1,94 | -2,26 | -1,91 | -1,80 | -2,09 | -1,59 |
| Topbp1 | -0,52 | -0,75 | -0,75 |  | -0,48 |  |
| Uvssa |  |  |  |  | -1,25 | -1,01 |
| Wrn | -1,23 | -1,10 | -1,21 | -1,32 | -1,59 | -1,06 |
| Xrcc4 | -1,01 | -1,17 | -1,14 | -1,16 | -0,55 | -0,92 |
| Xrcc5 | -1,50 | -1,12 | -1,52 | -1,32 | -0,88 | -0,95 |
| Xrcc6 | -0,52 | -0,40 | -0,40 |  |  |  |

Notably, after removal of *Bod1l*, “DNA repair” and related subterms were the predominant GO terms in all three cell states indicating that BOD1L action in SETD1A-C is focused on the DNA repair network (Fig. 6C). Notable downregulated genes after conditional mutagenesis of both *Bod1l* and *Setd1a* include mRNAs encoding proteins involved predominantly in homologous recombination repair, but also other pathways e.g. MLH1, ERCC6L2, FANCD2, RAD50, RAD51C, Werners syndrome helicase WRN, XRCC4 and 5, amongst others (Table 1). The promoters of DEGs decreased after removal of BOD1L also displayed increased H3K4me3 (Supplemental Fig. 6E).

As previously documented in cycling ESCs, loss of MLL2 promoted only modest changes in gene expression (Glaser et al. 2009; Denissov et al. 2014; Hu et al. 2017) with no concordance to changes after loss of SETD1A or BOD1L.

We then investigated the response to DNA damage by etoposide in the presence or absence of BOD1L (Fig. 7A). Etoposide provoked a strong upregulation of genes, including the TP53 transcriptional response, as expected. However this upregulation was muted by the absence of BOD1L (Fig. 7B). Notably the DNA damage caused by BOD1L deficiency induced a similar transcriptional response as etoposide (Fig.7C). We conclude that BOD1L is required for expression of many genes involved in DNA repair, including Homology Directed Repair. Additionally, the absence of BOD1L leads to the accumulation of DNA damage, which induces a damage response similar to the DNA damage response induced by etoposide.

**Figure 7.**
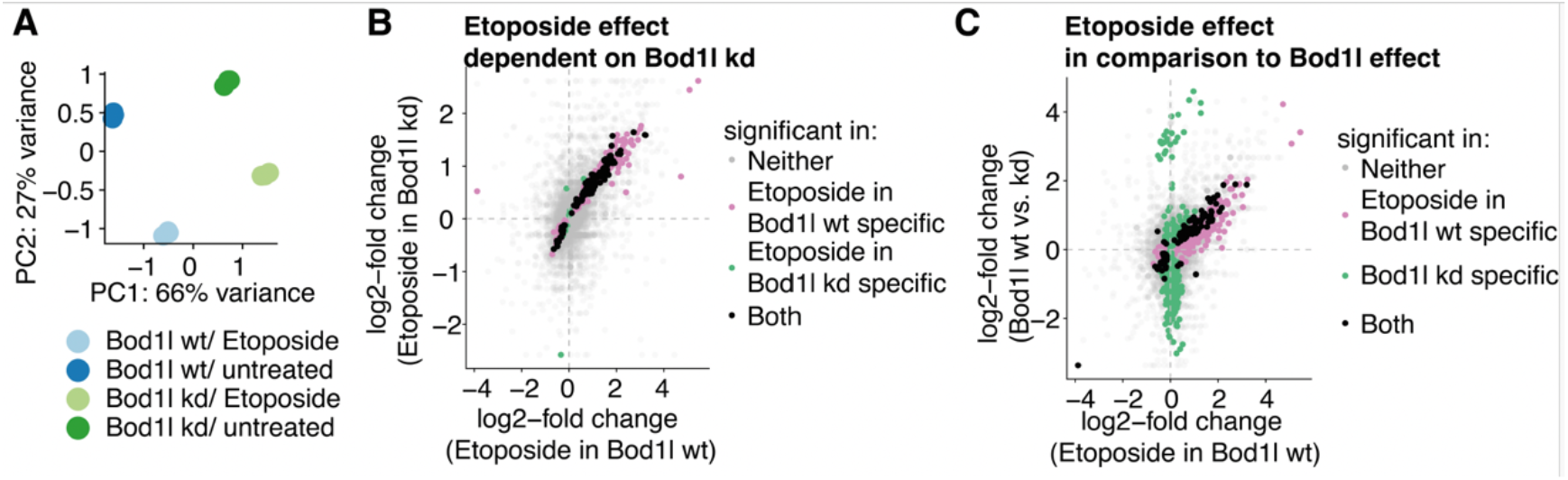
DNA damage responses induced by etoposide and the absence of BOD1L. **A.** Principal Component analysis of the transcriptomes dependent on *Bod1l* and etoposide treatment. **B.** Pairwise comparisons of gene expression dependent log2 fold changes from the differential analysis of etoposide induced gene expression dependent on Bod1l. Genes are highlighted when significant with an adjusted p-value < 0.05 within the stated condition. For visibility, axes were capped at the absolute maximum value of the genes significant in any samples or at a log2-fold change of 10. **C.** Pairwise comparisons of gene expression dependent log2 fold changes from the differential analysis of etoposide induced gene expression compared to gene expression changes without etoposide, but dependent on Bod1l. Genes are highlighted when significant with an adjusted p-value < 0.05 within the stated condition. For visibility, axes were capped at the absolute maximum value of the genes significant in any samples or at a log2-fold change of 10.

## Discussion

Of the six mammalian H3K4 methyltransferases, it was previously inferred that SETD1A is the pre-eminent methyltransferase in mammalian cells (Wu et al. 2008). Consistent with this pre-eminence, *Setd1a* knock-out in mice is required earlier in development than the knock-out of any of the other H3K4 methyltransferases (Ernst et al. 2004; Glaser et al. 2006; Bledau et al. 2014; Ashokkumar et al. 2020) and ubiquitous conditional mutagenesis of *Setd1a* in adult mice (using *Rosa26CreERT2*) provoked rapid death (Arndt et al. 2018). However, further investigations revealed that *Setd1a* is not required in certain circumstances including during oogenesis, early development before the epiblast (Bledau et al. 2014; Brici et al. 2017) and in hematopoietic stem cells (HSCs) where SETD1A is not required unless stress is imposed. Transcriptome profiling after loss of SETD1A revealed that HSCs fail under stress due to inadequate expression of many DNA damage repair genes (Arndt et al. 2018). Rather than playing a general role in gene expression as expected from the ubiquity of H3K4 methylation, SETD1A secures expression of genes involved in the DNA damage repair network, as well as genes involved in certain other central cellular functions including the mitochondria, cilia and Golgi. This selectivity is unexpected and resonates with the unexpected, more specialized, roles played by its paralog, SETD1B, which orchestrates a specific aspect of the gene expression program during oogenesis (Brici et al. 2017) and is required for appropriate expression of lineage specifying transcription factors during hematopoiesis (Schmidt et al. 2018). These observations raise the question: How do the two SET1Cs selectively sustain different gene expression programs?

Here we report that in ESCs the BOD1L subunit of SETD1A-C is required for the function of SETD1A in maintaining the expression of many DNA repair genes. Upon removal of either BOD1L or SETD1A, the expression of these DNA repair mRNAs decreases and unrepaired DNA damage accumulates leading to an inadequate DNA damage response and death. As shown by Grant Stewart and colleagues, BOD1L suppresses the degradation of stalled replication forks and interacts with RIF1 to enhance double strand break repair (DSBR) by non-homologous end joining (Higgs et al. 2015; Higgs et al. 2018; Bayley et al. 2022). Notably, SETD1A methyltransferase activity is required for DNA damage repair (Bayley et al. 2022). We suggest that in ESCs, the failure to repair DNA damage after loss of BOD1L is likely due to loss of both the direct action of BOD1L in DNA repair as well as the partial collapse of DNA repair gene expression due to defective SETD1A-C action. If BOD1L is exclusively involved in DNA damage repair without a contribution to the role of SETD1A-C in gene expression, then loss of BOD1L should lead to the elevated expression of DNA repair genes (Esposito and Francia 2026). However we observe decreased expression of many DNA repair genes, in particular *Xrcc4, Xrcc5* and *Rad51c*. πHence, we conclude that BOD1L is an intersectional node for DNA damage repair with two distinguishable functions; one in SETD1A-C to promote expression of various DNA repair genes and the other to protect the replication fork (Fig. 8). Concordantly, BOD1L is a prime target for ATR/ATM kinases and becomes highly phosphorylated in response to DNA damage (Matsuoka et al. 2007; Beli et al. 2012). The integration of two DNA repair functions in BOD1L likely conveys further implications for the DNA repair network. Whether the additional interactions of BOD1L with BPTF and SETD1B described here also contribute to DNA repair requires further investigation.

**Figure 8.**
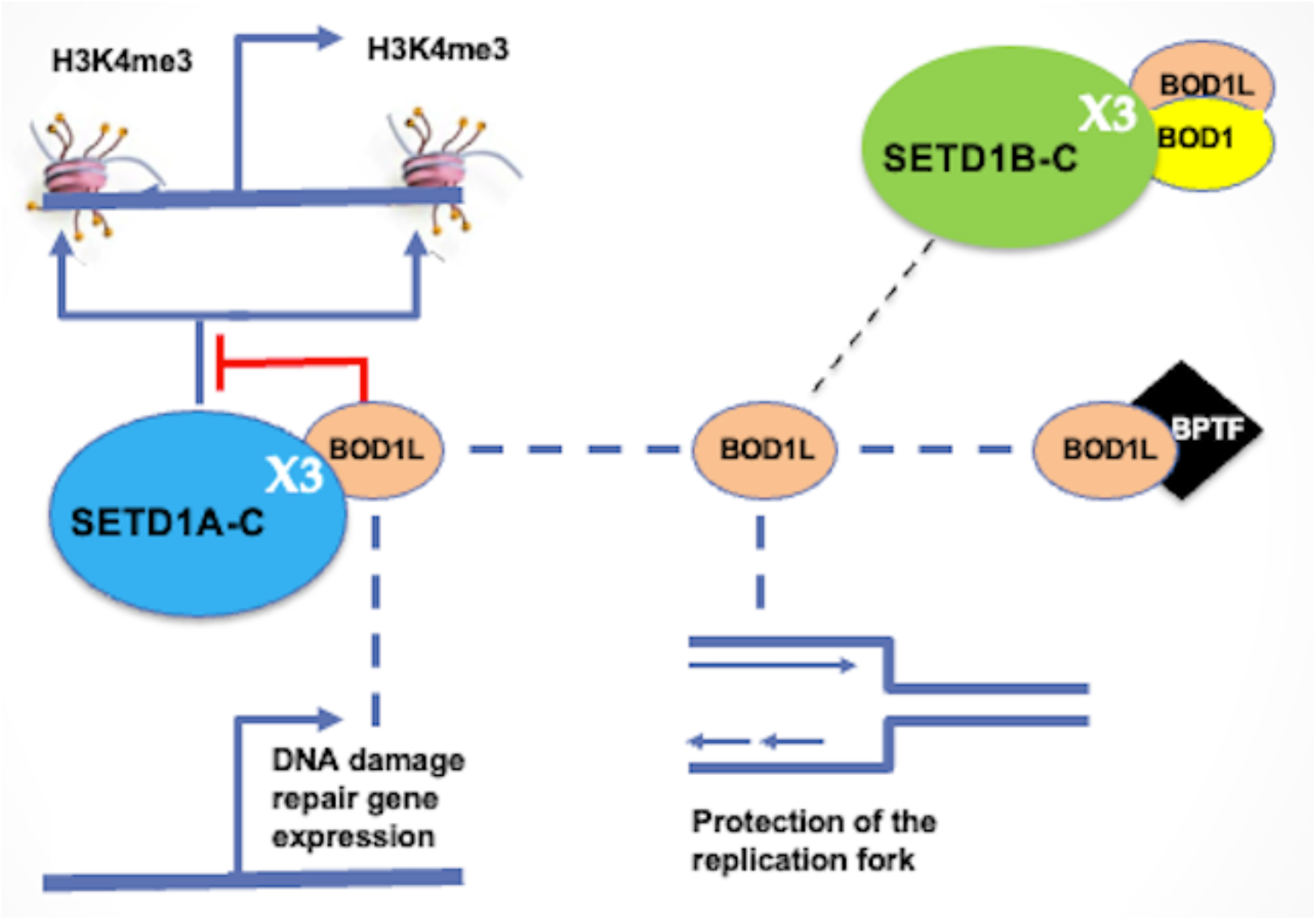
BOD1L interactions in ESCs. BOD1L participates in two aspects of the DNA damage response: (i) as a subunit of SETD1A-C to sustain the expression of many genes involved in DNA damage repair (see Table 1); and (ii) to protect the replication fork and enhance double strand break repair (DSBR) by non-homologous end joining (Higgs et al. 2015; Higgs et al. 2018; Bayley et al. 2022). BOD1L also interacts with BPTF and SETD1B. BOD1L binds to the X3 helix of SETD1A and restrains the trimethylation of H3K4 on the two nucleosomes that surround the start site of Pol II promoters.

Using a mouse leukemia cell line transformed by the MLL-AF9 translocation, Hoshii *et al* (Hoshii et al. 2018) also reported a relationship between SETD1A and DNA damage repair gene expression. Using a rescue assay with human *Setd1a* cDNA expression after knock-down of endogenous mouse *Setd1a*, a deletion analysis identified a requirement for human SETD1A amino acids 387 to 786, which they termed FLOS (functional location on SETD1A), as the region that conveys the rescue of the loss of DNA repair. Within FLOS, Hoshii *et al* defined four conserved elements termed FLOS 1 to 4. Probably FLOS1 is the same as X2, however the other FLOS assignments were not clearly specified. The X3 BOD1L interaction helix defined here starts at amino acid 786 so is excluded from the FLOS region. However, we note that in Figure 2C of Hoshii *et al*, 2018, the human cDNA 786-1026 deletion, which removes X3 but not FLOS, also failed to rescue. Although Hoshii *et al* did not pursue this observation at the time, we suggest that their results reveal the contribution of helix X3 and the BOD1L interaction for DNA repair, which also suggests that the role of BOD1L in DNA repair is likely to be present in many mammalian cell types.

Subsequent to the original bioRxiv posting of this manuscript containing the identification of helix X3 as the BOD1L interaction site, Hoshii *et al* published further observations relevant to this issue (Hoshii et al. 2024). In this later paper, they extended the FLOS region to add a fifth site to encompass the X3 interaction helix and report the interaction of BOD1L with this region including its’ relevance to DNA damage response in a leukemia cell line. Notably they did not observe elevated H3K4me3 when *Bod1l* was knocked out in this leukemia cell line.

Both BOD1L and BOD1 interact with the highly conserved X3 α-helix that lies in the middle of SETD1A and B. Our observations indicate that both combinations are possible; namely both BOD1L and BOD1 can bind to both SETD1A and B. However, it appears that SETD1A-BOD1L is the predominant species in ESCs (Figure 2) and HeLa cells (van Nuland et al. 2013). AlphaFold modelling of the interaction suggests that the Shg1 homology region masks the X3 α-helix (Supplemental Fig. S3). In some way, removal of Shg1/BOD1L binding to X3 promotes methylation by the highly conserved quintet of the WRAD (WDR5/RBBP5/ASH2L/DPY30) subcomplex, which is assembled on the SET domain at the C-terminus (Hsu et al. 2018; Qu et al. 2018; Worden et al. 2020). How does Shg1/BOD1L bound at the X3 helix regulate the SET domain? Various explanations remain possible including intramolecular interaction within Set1/SETD1A, intermolecular interaction within a Set1C/SETD1A-C homodimer or some other, as yet unidentified, X3 binding interaction. In this regard it is notable that binding of mono-ubiquitylated H2B by the SETD1C subunit, WDR82, is required for H3K4me3 (Wu et al. 2008) and WDR82 binds to the N-terminus of SETD1A (Hughes et al. 2023). Mechanistic explanations for the regulation of methylation by the C-terminal WRAD complex by the N-terminally distant binding of Shg1/BOD1L and WDR82 are lacking.

Encouraged by the AlphaFold predictions, we examined various putative BOD1 and BOD1L interaction partners including PP2A-5B (PP2A-B56), which is the 56-kDa beta subunit known to interact with BOD1 (Porter et al. 2013). The AlphaFold PP2A-5B prediction includes a prominent C-terminal α-helix, to which BOD1 binding would induce a conformational change (Supplemental Fig. S4).

All eukaryotes examined so far have at least one Set1 H3K4 methyltransferase and H3K4 methylation is a universal characteristic of active chromatin, including the ubiquity of H3K4me3 on nucleosomes surrounding active Pol II promoters (Talbert and Henikoff 2021). These discoveries promoted investigations of H3K4me3 as an epitope in promoter architecture and the mechanics of transcriptional initiation. The identification of the TAF3 subunit of TFIID as an H3K4me3 binder secured these investigations (Vermeulen et al. 2007; Lauberth et al. 2013), which has been recently extended by the identification of a role for H3K4me3 in promoter-proximal pause-release of RNA polymerase II (Hu et al. 2023; Wang et al. 2023). (Hughes et al. 2023).

Alternative, epigenetic, explanations for the ubiquity of H3K4 methylation arose from the discoveries that the genetic opposition between TrxG and PcG action related to competition for the methylation status of H3, with H3K4 methylation mediated by TrxG action and mutually exclusive H3K27me3 mediated by PcG action (Klymenko and Muller 2004; Kim et al. 2013a). Similarly, in mammals, H3K4me3 repels a variety of repressive chromatin processes including DNA methylation (Ooi et al. 2007; Eberl et al. 2013; Stewart et al. 2016; Hanna et al. 2018). Furthermore, observations in yeast and flies indicated that gene expression does not rely on H3K4me3 (Lenstra et al. 2011; Hodl and Basler 2012; Margaritis et al. 2012; Weiner et al. 2012). At least in yeast, H3K4me3 appears to be a consequence of transcription rather than a prerequisite (Soares et al. 2017; Choudhury et al. 2019)

Although explanations based on both epigenetic and transcriptional mechanics and are compatible, indeed complementary, our observation that loss of SETD1A and BOD1L provoke similar changes in gene expression but converse changes in H3K4me3 indicates that H3K4me3 is peripheral to the action of SETD1A-C in sustaining gene expression in ESCs. This conclusion tends to emphasize the primary importance of H3K4 methylation in epigenetics and the reinforcing mechanisms that establish chromatin status utilizing feed-forward loops (Jaenisch and Bird 2003; Talbert and Henikoff 2021)

Amongst the candidate interactors identified by AP-MS, we confirmed the BOD1L interaction with BPTF. We also note multiple identifications of SET1-C with (i) several of the MLL4 complex subunits (MLL4/KMT2D, UTX/KDM6A, PAXIP1, NCOA6); (ii) the entire cleavage polyadenylation complex (CSTF, CPSF, WDR33); and (iii) ESRRB (Figs. 2-4). Examination of these concurrences could be fruitful. In particular, low affinity relationships between SET1-Cs and MLL-Cs is suggestive of redundancies and overlapping complexities amongst the H3K4 methyltransferases, which would extend previously reported functional overlaps for MLL1 and 2 (Denissov et al. 2014; Chen et al. 2017) and MLL3 and 4 (Dorighi et al. 2017; Ashokkumar et al. 2020; Kubo et al. 2024; Wang et al. 2025).

## Material and Methods

### Targeting constructs, gene targeting and BAC transgenes

The *Bod1l* targeting constructs and BAC transgenes were generated using recombineering (Fu et al. 2010; Ciotta et al. 2011; Hofemeister et al. 2011) using BACs (CH29-573N17, CH29-558P19, RP23-320D8, RP23-450O9, RP23-117I12, bMQ298H3) containing *Bod1l, Bod1, Cxxc1, Bptf*, *Mll2* and *Mll4* genes respectively, which were purchased from CHORI (Children’s Hospital Oakland Research Institute; http://bacpac.chori.org) or Geneservice (http://www.geneservice.co.uk). Both 2xTy1-X3 coding regions were ordered as two overlapping 130mer oligonucleotides, filled in with Klenow and ligated into pCAG-IRESpuro.

### ESC culture

Culture and targeting of mouse ES cells were performed as previously described (Anastassiadis et al. 2013). ESC2i and ESCFCS were cultured as described (Marks et al. 2012). ESCs differentiated towards epiblast-like stem cells (EpiSCs) by culture without LIF in the presence of Activin A and FGF2 (Hayashi et al. 2011). Conversion of floxed alleles, denoted F/F, to deleted versions, denoted FC/FC, was achieved by addition of 500 nM 4-hydroxy tamoxifen to the culture for 48 hours, except for the etoposide experiment when it was 72 hours, followed by 0.1μM etoposide for 6 hours before harvesting.

### AP-MS

GFP/Venus-tagged protein complexes were isolated by immunoaffinity chromatography using a fully automated liquid-handling platform as described (Hubner et al. 2010). LC-MS/MS analysis was performed by using a mass spectrometer (Q Exactive; Thermo Scientific). The Q-Exactive was operated using Xcalibur 2.2 in the data-dependent mode to automatically switch between MS and MS/MS acquisition as described (Michalski et al. 2011; Kelstrup et al. 2012). Raw data files were processed using the MaxQuant software (v1.2.6.20; https://www.maxquant.org/) as described previously (Cox et al. 2011). Parent ion (MS) and fragment (MS2) spectra were searched against the UniProt species-specific fasta files from the February 2012 release. Label-free quantification was performed with the Perseus software (https://www.maxquant.org/perseus/) and the MaxQuant-based program QUBICvalidator as described previously (Hubner et al. 2010). Proteins with more than 2 unique peptides in all 3 triplicate samples were further analyzed. Results were then plotted by using the open source statistical software R (https://www.Rproject.org). The Volcano plots of Figures 2 - 4 were generated using MS-Volcano (Singh et al. 2016).

### Antibodies, immunohistochemistry, immunoprecipitation and Western analysis

ES cells were homogenized in buffer E (20 mM HEPES, 350 mM NaCl, 10% Glycerol, 0.1% Tween 20, 1 μg/ml pepstatin A, 0.5 μg/ml leupeptin, 2 μg/ml aprotinin, 1 mM PMSF) and snap-frozen three times for protein extraction. Protein extracts were fractionated by SDS-PAGE (3-8% Tris-acetate gel for Setd1 proteins, 12% Tris-Glycine gel for histone proteins), transferred to PVDF membrane and probed with primary antibodies: GFP (Denissov et al. 2014), SETD1A (1:500, A300-289 Bethyl), SETD1B (1:500, A302-281A Bethyl), CTR9 (1:500, A301-395A Bethyl), BPTF (1:50, HPA029069 Sigma), SNF2L (39952 Biobyt), MLL1 (A300-086A Bethyl, (!!! INVALID CITATION !!! (Goveas et al. 2021)), MLL2 (Glaser et al. 2006), H3 (1:3500, ab1791 Abcam), H3K4me1 (1:2500, ab889 Abcam), H3K4me2 (1:2500, ab7766 Abcam), H3K4me3 (1:1000, 07-473, Millipore), H3K27me3 (1:2500, 07-449 Millipore), H2A.X phosphoS139 (Cell Signaling 2577), ATM phosphoS1981 (ThermoFischer MA1-2020 10H11). The Ty1 BB2 monoclonal antibody (Brookman et al. 1995) was an IgG enriched fraction.

### RNA-seq and ChIP-seq

RNA-seq and ChIP-seq were performed as described (Denissov et al. 2014). Single-end bulk RNA-seq reads from 36 samples were mapped to the mouse reference genome GRCm38 using GSNAP (version 2017-03-17) and paired-end reads from 12 samples were aligned to GRCm39 using Hisat2 (version 2.2.1). Gene expression was quantified using featureCounts (version 1.5.2) for the single-end data and StringTie (version 2.2.1) for the paired-end data. Differential expression analysis was performed with the DESeq2 package (version 1.48.2) (Love et al. 2014) within the R programming environment (version 4.5.1.) (http://www.R-project.org/). Comparisons of log2-fold changes were visualised with an axis cap at the absolute maximal value of a significant gene or a log2-fold change of 10. Mitochondrial genes were filtered out. An adjusted p-value of ≤0.05 was used as the significance threshold for the identification of differentially expressed genes. Heatmaps to show differentially expressed genes were generated on count data that were normalised and norm transformed using the DESeq2 package, and depicted using “pheatmap” with scaling by row. GO term analysis was performed with the gprofiler2 package (version 0.2.3) within the R programming environment (version 4.5.1.) (http://www.R-project.org/) using reactome.db_(version 1.92.0), ReactomePA (version 1.52.0). GO terms were considered with a significance threshold of 0.05 and the top 10 are visualised.

Single-end H3K4me3 ChIP-seq data from control and *Bod1l* knock-out cells were processed in separate runs using the ENCODE-DCC chip-seq-pipeline2 (version 2.2.2)(Hitz et al. 2023). Each condition comprised two biological replicates. Three mock-IP libraries were pooled and used as a common control for peak calling. Reads were mapped to the mouse GRCm38/mm10 reference genome using Bowtie2. Narrow H3K4me3 peaks were called using MACS2, and the optimal reproducible peak sets were used for downstream analysis. Active promoters were defined from the control samples as regions ±1 kb around gene TSS that overlapped an H3K4me3 peak. Promoters of genes with a mean RNA-seq expression below 1 transcript per million (TPM) across the control and *Bod1l* knock-out ESC2i samples were excluded. For signal normalisation, H3K4me3 peaks from the control and *Bod1l* knock-out samples were merged into a common non-overlapping peak set. Reads from each filtered, de-duplicated BAM file were counted within these regions using featureCounts (version 2.1.1). Regions with at least 10 reads in at least two libraries were retained. TMM normalisation factors were calculated using calcNormFactors in edgeR (version 4.4.2) with the “method” parameter set to “TMM”. For each sample, the effective peak-set library size was calculated as the total read count within the retained regions multiplied by the TMM normalisation factor. The size factor was divided by 10^6^, and its reciprocal was supplied to bamCoverage (version 3.5.6) in deepTools to generate TMM-normalised bigWig files.

## Supporting information

Response to Reviewers

## Acknowledgements

AFS and CC were primarily funded by the EU Integrated project, SyBoSS https://cordis.europa.eu/project/id/242129/reporting/es. The Novo Nordisk Foundation Center for Protein Research is financially supported by the Novo Nordisk Foundation (NNF14CC0001). Computational resources were provided by both the BMBF project scads.ai and the e-INFRA CZ project (ID:90140), supported by the Ministry of Education, Youth and Sports of the Czech Republic. ARP was supported by the Mildred Scheel Early Career Center Dresden P2, funded by the German Cancer Aid. We thank the Dresden concept Genome Center and Andreas Petzold for excellent sequencing services, Mandy Obst for technical assistance and Michelle Meredyth for critical review of the manuscript.

## Author contributions

GC, AG, DCT and JF did the molecular biology (including recombineering, AP, immunoprecipitation, Westerns). WKC and CC did the AP-MS and early MS analysis. GC, AG, DCT, AK, NL, KA did the ESC work. RA did the Shg1/BOD and SET1 alignments. AA-F and MS did the AlphaFold analyses. SS, LG, RC, DS, ARP and AFS did the other computational analyses. KA, ARP, AFS wrote the paper with input from all co-authors.

## Notes

https://www.ncbi.nlm.nih.gov/geo/query/acc.cgi?acc=GSE319676

https://www.ncbi.nlm.nih.gov/geo/query/acc.cgi?acc=GSE319677

### Competing Interest Statement

The authors have declared no competing interest.

### Summary of Updates

The manuscript has been revised in response to three reviewers as part of the Review Commons process. The principle addition involves Supplemental Figure 6E, which shows that elevated H3K4me3 after Bod1l conditional mutagenesis is not only generally observed on active promoters but also on the promoters of DNA repair genes that show decreased expression. In the text, certain errors have been removed and expression improved in places.

## References

Anastassiadis K, Schnutgen F, von Melchner H, Stewart AF. 2013. Gene targeting and site-specific recombination in mouse ES cells. Methods Enzymol 533: 133–155.

Ardehali MB, Mei A, Zobeck KL, Caron M, Lis JT, Kusch T. 2011. Drosophila Set1 is the major histone H3 lysine 4 trimethyltransferase with role in transcription. EMBO J 30: 2817–2828.

Arndt K, Kranz A, Fohgrub J, Jolly A, Bledau AS, Di Virgilio M, Lesche M, Dahl A, Hofer T, Stewart AF et al. 2018. SETD1A protects HSCs from activation-induced functional decline in vivo. Blood 131: 1311–1324.

Ashokkumar D, Zhang Q, Much C, Bledau AS, Naumann R, Alexopoulou D, Dahl A, Goveas N, Fu J, Anastassiadis K et al. 2020. MLL4 is required after implantation, whereas MLL3 becomes essential during late gestation. Development 147.

Bayley R, Borel V, Moss RJ, Sweatman E, Ruis P, Ormrod A, Goula A, Mottram RMA, Stanage T, Hewitt G et al. 2022. H3K4 methylation by SETD1A/BOD1L facilitates RIF1-dependent NHEJ. Mol Cell 82: 1924–1939 e1910.

Beli P, Lukashchuk N, Wagner SA, Weinert BT, Olsen JV, Baskcomb L, Mann M, Jackson SP, Choudhary C. 2012. Proteomic investigations reveal a role for RNA processing factor THRAP3 in the DNA damage response. Mol Cell 46: 212–225.

Bledau AS, Schmidt K, Neumann K, Hill U, Ciotta G, Gupta A, Torres DC, Fu J, Kranz A, Stewart AF et al. 2014. The H3K4 methyltransferase Setd1a is first required at the epiblast stage, whereas Setd1b becomes essential after gastrulation. Development 141: 1022–1035.

Brici D, Zhang Q, Reinhardt S, Dahl A, Hartmann H, Schmidt K, Goveas N, Huang J, Gahurova L, Kelsey G et al. 2017. Setd1b, encoding a histone 3 lysine 4 methyltransferase, is a maternal effect gene required for the oogenic gene expression program. Development 144: 2606–2617.

Brookman JL, Stott AJ, Cheeseman PJ, Burns NR, Adams SE, Kingsman AJ, Gull K. 1995. An immunological analysis of Ty1 virus-like particle structure. Virology 207: 59–67.

Chen Y, Anastassiadis K, Kranz A, Stewart AF, Arndt K, Waskow C, Yokoyama A, Jones K, Neff T, Lee Y et al. 2017. MLL2, Not MLL1, Plays a Major Role in Sustaining MLL-Rearranged Acute Myeloid Leukemia. Cancer Cell 31: 755–770 e756.

Choudhary C, Mann M. 2010. Decoding signalling networks by mass spectrometry-based proteomics. Nat Rev Mol Cell Biol 11: 427–439.

Choudhury R, Singh S, Arumugam S, Roguev A, Stewart AF. 2019. The Set1 complex is dimeric and acts with Jhd2 demethylation to convey symmetrical H3K4 trimethylation. Genes Dev 33: 550–564.

Ciotta G, Hofemeister H, Maresca M, Fu J, Sarov M, Anastassiadis K, Stewart AF. 2011. Recombineering BAC transgenes for protein tagging. Methods 53: 113–119.

Cox J, Neuhauser N, Michalski A, Scheltema RA, Olsen JV, Mann M. 2011. Andromeda: a peptide search engine integrated into the MaxQuant environment. J Proteome Res 10: 1794–1805.

Dehe PM, Dichtl B, Schaft D, Roguev A, Pamblanco M, Lebrun R, Rodriguez-Gil A, Mkandawire M, Landsberg K, Shevchenko A et al. 2006. Protein interactions within the Set1 complex and their roles in the regulation of histone 3 lysine 4 methylation. J Biol Chem 281: 35404–35412.

Denissov S, Hofemeister H, Marks H, Kranz A, Ciotta G, Singh S, Anastassiadis K, Stunnenberg HG, Stewart AF. 2014. Mll2 is required for H3K4 trimethylation on bivalent promoters in embryonic stem cells, whereas Mll1 is redundant. Development 141: 526–537.

Dorighi KM, Swigut T, Henriques T, Bhanu NV, Scruggs BS, Nady N, Still CD, 2nd, Garcia BA, Adelman K, Wysocka J. 2017. Mll3 and Mll4 Facilitate Enhancer RNA Synthesis and Transcription from Promoters Independently of H3K4 Monomethylation. Mol Cell 66: 568–576 e564.

Eberl HC, Spruijt CG, Kelstrup CD, Vermeulen M, Mann M. 2013. A map of general and specialized chromatin readers in mouse tissues generated by label-free interaction proteomics. Mol Cell 49: 368–378.

Ernst P, Fisher JK, Avery W, Wade S, Foy D, Korsmeyer SJ. 2004. Definitive hematopoiesis requires the mixed-lineage leukemia gene. Dev Cell 6: 437–443.

Ernst P, Vakoc CR. 2012. WRAD: enabler of the SET1-family of H3K4 methyltransferases. Brief Funct Genomics 11: 217–226.

Esposito F, Francia S. 2026. DNA double-strand break response at a glance. J Cell Sci 139.

Feil R, Wagner J, Metzger D, Chambon P. 1997. Regulation of Cre recombinase activity by mutated estrogen receptor ligand-binding domains. Biochem Biophys Res Commun 237: 752–757.

Fu J, Teucher M, Anastassiadis K, Skarnes W, Stewart AF. 2010. A recombineering pipeline to make conditional targeting constructs. Methods Enzymol 477: 125–144.

Gambetta MC, Oktaba K, Muller J. 2009. Essential role of the glycosyltransferase sxc/Ogt in polycomb repression. Science 325: 93–96.

Glaser S, Lubitz S, Loveland KL, Ohbo K, Robb L, Schwenk F, Seibler J, Roellig D, Kranz A, Anastassiadis K et al. 2009. The histone 3 lysine 4 methyltransferase, Mll2, is only required briefly in development and spermatogenesis. Epigenetics Chromatin 2: 5.

Glaser S, Schaft J, Lubitz S, Vintersten K, van der Hoeven F, Tufteland KR, Aasland R, Anastassiadis K, Ang SL, Stewart AF. 2006. Multiple epigenetic maintenance factors implicated by the loss of Mll2 in mouse development. Development 133: 1423–1432.

Hanna CW, Taudt A, Huang J, Gahurova L, Kranz A, Andrews S, Dean W, Stewart AF, Colome-Tatche M, Kelsey G. 2018. MLL2 conveys transcription-independent H3K4 trimethylation in oocytes. Nat Struct Mol Biol 25: 73–82.

Hayashi K, Ohta H, Kurimoto K, Aramaki S, Saitou M. 2011. Reconstitution of the mouse germ cell specification pathway in culture by pluripotent stem cells. Cell 146: 519–532.

Higgs MR, Reynolds JJ, Winczura A, Blackford AN, Borel V, Miller ES, Zlatanou A, Nieminuszczy J, Ryan EL, Davies NJ et al. 2015. BOD1L Is Required to Suppress Deleterious Resection of Stressed Replication Forks. Mol Cell 59: 462–477.

Higgs MR, Sato K, Reynolds JJ, Begum S, Bayley R, Goula A, Vernet A, Paquin KL, Skalnik DG, Kobayashi W et al. 2018. Histone Methylation by SETD1A Protects Nascent DNA through the Nucleosome Chaperone Activity of FANCD2. Mol Cell 71: 25–41 e26.

Hitz BC, Lee JW, Jolanki O, Kagda MS, Graham K, Sud P, Gabdank I, Strattan JS, Sloan CA, Dreszer T et al. 2023. The ENCODE Uniform Analysis Pipelines. Res Sq.

Hodl M, Basler K. 2012. Transcription in the absence of histone H3.2 and H3K4 methylation. Curr Biol 22: 2253–2257.

Hofemeister H, Ciotta G, Fu J, Seibert PM, Schulz A, Maresca M, Sarov M, Anastassiadis K, Stewart AF. 2011. Recombineering, transfection, Western, IP and ChIP methods for protein tagging via gene targeting or BAC transgenesis. Methods 53: 437–452.

Hoshii T, Cifani P, Feng Z, Huang CH, Koche R, Chen CW, Delaney CD, Lowe SW, Kentsis A, Armstrong SA. 2018. A Non-catalytic Function of SETD1A Regulates Cyclin K and the DNA Damage Response. Cell 172: 1007–1021 e1017.

Hoshii T, Kikuchi S, Kujirai T, Masuda T, Ito T, Yasuda S, Matsumoto M, Rahmutulla B, Fukuyo M, Murata T et al. 2024. BOD1L mediates chromatin binding and non-canonical function of H3K4 methyltransferase SETD1A. Nucleic Acids Res 52: 9463–9480.

Howe FS, Fischl H, Murray SC, Mellor J. 2017. Is H3K4me3 instructive for transcription activation? Bioessays 39: 1–12.

Hsu PL, Li H, Lau HT, Leonen C, Dhall A, Ong SE, Chatterjee C, Zheng N. 2018. Crystal Structure of the COMPASS H3K4 Methyltransferase Catalytic Module. Cell 174: 1106–1116 e1109.

Hu D, Gao X, Cao K, Morgan MA, Mas G, Smith ER, Volk AG, Bartom ET, Crispino JD, Di Croce L et al. 2017. Not All H3K4 Methylations Are Created Equal: Mll2/COMPASS Dependency in Primordial Germ Cell Specification. Mol Cell 65: 460–475 e466.

Hu S, Song A, Peng L, Tang N, Qiao Z, Wang Z, Lan F, Chen FX. 2023. H3K4me2/3 modulate the stability of RNA polymerase II pausing. Cell Res 33: 403–406.

Hubner NC, Bird AW, Cox J, Splettstoesser B, Bandilla P, Poser I, Hyman A, Mann M. 2010. Quantitative proteomics combined with BAC TransgeneOmics reveals in vivo protein interactions. J Cell Biol 189: 739–754.

Hughes AL, Szczurek AT, Kelley JR, Lastuvkova A, Turberfield AH, Dimitrova E, Blackledge NP, Klose RJ. 2023. A CpG island-encoded mechanism protects genes from premature transcription termination. Nat Commun 14: 726.

Jaenisch R, Bird A. 2003. Epigenetic regulation of gene expression: how the genome integrates intrinsic and environmental signals. Nat Genet 33 Suppl: 245–254.

Kelstrup CD, Young C, Lavallee R, Nielsen ML, Olsen JV. 2012. Optimized fast and sensitive acquisition methods for shotgun proteomics on a quadrupole orbitrap mass spectrometer. J Proteome Res 11: 3487–3497.

Kim DH, Tang Z, Shimada M, Fierz B, Houck-Loomis B, Bar-Dagen M, Lee S, Lee SK, Muir TW, Roeder RG et al. 2013a. Histone H3K27 trimethylation inhibits H3 binding and function of SET1-like H3K4 methyltransferase complexes. Mol Cell Biol 33: 4936–4946.

Kim J, Kim JA, McGinty RK, Nguyen UT, Muir TW, Allis CD, Roeder RG. 2013b. The n-SET domain of Set1 regulates H2B ubiquitylation-dependent H3K4 methylation. Mol Cell 49: 1121–1133.

Klymenko T, Muller J. 2004. The histone methyltransferases Trithorax and Ash1 prevent transcriptional silencing by Polycomb group proteins. EMBO Rep 5: 373–377.

Kranz A, Anastassiadis K. 2020. The role of SETD1A and SETD1B in development and disease. Biochim Biophys Acta Gene Regul Mech 1863: 194578.

Krogan NJ, Dover J, Khorrami S, Greenblatt JF, Schneider J, Johnston M, Shilatifard A. 2002. COMPASS, a histone H3 (Lysine 4) methyltransferase required for telomeric silencing of gene expression. J Biol Chem 277: 10753–10755.

Kubo N, Chen PB, Hu R, Ye Z, Sasaki H, Ren B. 2024. H3K4me1 facilitates promoter-enhancer interactions and gene activation during embryonic stem cell differentiation. Mol Cell 84: 1742–1752 e1745.

Lauberth SM, Nakayama T, Wu X, Ferris AL, Tang Z, Hughes SH, Roeder RG. 2013. H3K4me3 interactions with TAF3 regulate preinitiation complex assembly and selective gene activation. Cell 152: 1021–1036.

Lee JH, Skalnik DG. 2005. CpG-binding protein (CXXC finger protein 1) is a component of the mammalian Set1 histone H3-Lys4 methyltransferase complex, the analogue of the yeast Set1/COMPASS complex. J Biol Chem 280: 41725–41731.

Lee JH, Skalnik DG. 2008. Wdr82 is a C-terminal domain-binding protein that recruits the Setd1A Histone H3-Lys4 methyltransferase complex to transcription start sites of transcribed human genes. Mol Cell Biol 28: 609–618.

Lee JH, Tate CM, You JS, Skalnik DG. 2007. Identification and characterization of the human Set1B histone H3-Lys4 methyltransferase complex. J Biol Chem 282: 13419–13428.

Lenstra TL, Benschop JJ, Kim T, Schulze JM, Brabers NA, Margaritis T, van de Pasch LA, van Heesch SA, Brok MO, Groot Koerkamp MJ et al. 2011. The specificity and topology of chromatin interaction pathways in yeast. Mol Cell 42: 536–549.

Li H, Ilin S, Wang W, Duncan EM, Wysocka J, Allis CD, Patel DJ. 2006. Molecular basis for site-specific read-out of histone H3K4me3 by the BPTF PHD finger of NURF. Nature 442: 91–95.

Logie C, Stewart AF. 1995. Ligand-regulated site-specific recombination. Proc Natl Acad Sci U S A 92: 5940–5944.

Love MI, Huber W, Anders S. 2014. Moderated estimation of fold change and dispersion for RNA-seq data with DESeq2. Genome Biol 15: 550.

Margaritis T, Oreal V, Brabers N, Maestroni L, Vitaliano-Prunier A, Benschop JJ, van Hooff S, van Leenen D, Dargemont C, Geli V et al. 2012. Two distinct repressive mechanisms for histone 3 lysine 4 methylation through promoting 3’-end antisense transcription. PLoS Genet 8: e1002952.

Marks H, Kalkan T, Menafra R, Denissov S, Jones K, Hofemeister H, Nichols J, Kranz A, Stewart AF, Smith A et al. 2012. The transcriptional and epigenomic foundations of ground state pluripotency. Cell 149: 590–604.

Matsuoka S, Ballif BA, Smogorzewska A, McDonald ER, 3rd, Hurov KE, Luo J, Bakalarski CE, Zhao Z, Solimini N, Lerenthal Y et al. 2007. ATM and ATR substrate analysis reveals extensive protein networks responsive to DNA damage. Science 316: 1160–1166.

Michalski A, Damoc E, Hauschild JP, Lange O, Wieghaus A, Makarov A, Nagaraj N, Cox J, Mann M, Horning S. 2011. Mass spectrometry-based proteomics using Q Exactive, a high-performance benchtop quadrupole Orbitrap mass spectrometer. Mol Cell Proteomics 10: M111 011015.

Miller T, Krogan NJ, Dover J, Erdjument-Bromage H, Tempst P, Johnston M, Greenblatt JF, Shilatifard A. 2001. COMPASS: a complex of proteins associated with a trithorax-related SET domain protein. Proc Natl Acad Sci U S A 98: 12902–12907.

Mohan M, Herz HM, Smith ER, Zhang Y, Jackson J, Washburn MP, Florens L, Eissenberg JC, Shilatifard A. 2011. The COMPASS family of H3K4 methylases in Drosophila. Mol Cell Biol 31: 4310–4318.

Morillon A, Karabetsou N, Nair A, Mellor J. 2005. Dynamic lysine methylation on histone H3 defines the regulatory phase of gene transcription. Mol Cell 18: 723–734.

Olivieri M, Cho T, Alvarez-Quilon A, Li K, Schellenberg MJ, Zimmermann M, Hustedt N, Rossi SE, Adam S, Melo H et al. 2020. A Genetic Map of the Response to DNA Damage in Human Cells. Cell 182: 481–496 e421.

Ooi SK, Qiu C, Bernstein E, Li K, Jia D, Yang Z, Erdjument-Bromage H, Tempst P, Lin SP, Allis CD et al. 2007. DNMT3L connects unmethylated lysine 4 of histone H3 to de novo methylation of DNA. Nature 448: 714–717.

Piunti A, Shilatifard A. 2016. Epigenetic balance of gene expression by Polycomb and COMPASS families. Science 352: aad9780.

Porter IM, McClelland SE, Khoudoli GA, Hunter CJ, Andersen JS, McAinsh AD, Blow JJ, Swedlow JR. 2007. Bod1, a novel kinetochore protein required for chromosome biorientation. J Cell Biol 179: 187–197.

Porter IM, Schleicher K, Porter M, Swedlow JR. 2013. Bod1 regulates protein phosphatase 2A at mitotic kinetochores. Nat Commun 4: 2677.

Qu Q, Takahashi YH, Yang Y, Hu H, Zhang Y, Brunzelle JS, Couture JF, Shilatifard A, Skiniotis G. 2018. Structure and Conformational Dynamics of a COMPASS Histone H3K4 Methyltransferase Complex. Cell 174: 1117–1126 e1112.

Roguev A, Schaft D, Shevchenko A, Aasland R, Shevchenko A, Stewart AF. 2003. High conservation of the Set1/Rad6 axis of histone 3 lysine 4 methylation in budding and fission yeasts. J Biol Chem 278: 8487–8493.

Roguev A, Schaft D, Shevchenko A, Pijnappel WW, Wilm M, Aasland R, Stewart AF. 2001. The Saccharomyces cerevisiae Set1 complex includes an Ash2 homologue and methylates histone 3 lysine 4. EMBO J 20: 7137–7148.

Schleicher K, Porter M, Ten Have S, Sundaramoorthy R, Porter IM, Swedlow JR. 2017. The Ndc80 complex targets Bod1 to human mitotic kinetochores. Open Biol 7.

Schmidt K, Zhang Q, Tasdogan A, Petzold A, Dahl A, Arneth BM, Slany R, Fehling HJ, Kranz A, Stewart AF et al. 2018. The H3K4 methyltransferase Setd1b is essential for hematopoietic stem and progenitor cell homeostasis in mice. Elife 7.

Schneider J, Wood A, Lee JS, Schuster R, Dueker J, Maguire C, Swanson SK, Florens L, Washburn MP, Shilatifard A. 2005. Molecular regulation of histone H3 trimethylation by COMPASS and the regulation of gene expression. Mol Cell 19: 849–856.

Schwenk F, Kuhn R, Angrand PO, Rajewsky K, Stewart AF. 1998. Temporally and spatially regulated somatic mutagenesis in mice. Nucleic Acids Res 26: 1427–1432.

Seibler J, Zevnik B, Kuter-Luks B, Andreas S, Kern H, Hennek T, Rode A, Heimann C, Faust N, Kauselmann G et al. 2003. Rapid generation of inducible mouse mutants. Nucleic Acids Res 31: e12.

Shevchenko A, Schaft D, Roguev A, Pijnappel WW, Stewart AF, Shevchenko A. 2002. Deciphering protein complexes and protein interaction networks by tandem affinity purification and mass spectrometry: analytical perspective. Mol Cell Proteomics 1: 204–212.

Shi X, Kachirskaia I, Walter KL, Kuo JH, Lake A, Davrazou F, Chan SM, Martin DG, Fingerman IM, Briggs SD et al. 2007. Proteome-wide analysis in Saccharomyces cerevisiae identifies several PHD fingers as novel direct and selective binding modules of histone H3 methylated at either lysine 4 or lysine 36. J Biol Chem 282: 2450–2455.

Singh S, Hein MY, Stewart AF. 2016. msVolcano: A flexible web application for visualizing quantitative proteomics data. Proteomics 16: 2491–2494.

Soares LM, He PC, Chun Y, Suh H, Kim T, Buratowski S. 2017. Determinants of Histone H3K4 Methylation Patterns. Mol Cell 68: 773–785 e776.

Stewart KR, Veselovska L, Kelsey G. 2016. Establishment and functions of DNA methylation in the germline. Epigenomics 8: 1399–1413.

Talbert PB, Henikoff S. 2021. The Yin and Yang of Histone Marks in Transcription. Annu Rev Genomics Hum Genet 22: 147–170.

Tsukiyama T, Wu C. 1995. Purification and properties of an ATP-dependent nucleosome remodeling factor. Cell 83: 1011–1020.

van Nuland R, Smits AH, Pallaki P, Jansen PW, Vermeulen M, Timmers HT. 2013. Quantitative dissection and stoichiometry determination of the human SET1/MLL histone methyltransferase complexes. Mol Cell Biol 33: 2067–2077.

Vermeulen M, Mulder KW, Denissov S, Pijnappel WW, van Schaik FM, Varier RA, Baltissen MP, Stunnenberg HG, Mann M, Timmers HT. 2007. Selective anchoring of TFIID to nucleosomes by trimethylation of histone H3 lysine 4. Cell 131: 58–69.

Wang H, Fan Z, Shliaha PV, Miele M, Hendrickson RC, Jiang X, Helin K. 2023. H3K4me3 regulates RNA polymerase II promoter-proximal pause-release. Nature 615: 339–348.

Wang N, Pachai MR, Li D, Lee CJ, Warda S, Khudoynazarova MN, Cho WH, Xie G, Shah SR, Yao L et al. 2025. Loss of Kmt2c or Kmt2d primes urothelium for tumorigenesis and redistributes KMT2A-menin to bivalent promoters. Nat Genet 57: 165–179.

Weiner A, Chen HV, Liu CL, Rahat A, Klien A, Soares L, Gudipati M, Pfeffner J, Regev A, Buratowski S et al. 2012. Systematic dissection of roles for chromatin regulators in a yeast stress response. PLoS Biol 10: e1001369.

Worden EJ, Zhang X, Wolberger C. 2020. Structural basis for COMPASS recognition of an H2B-ubiquitinated nucleosome. Elife 9.

Wu M, Wang PF, Lee JS, Martin-Brown S, Florens L, Washburn M, Shilatifard A. 2008. Molecular regulation of H3K4 trimethylation by Wdr82, a component of human Set1/COMPASS. Mol Cell Biol 28: 7337–7344.

Wysocka J, Myers MP, Laherty CD, Eisenman RN, Herr W. 2003. Human Sin3 deacetylase and trithorax-related Set1/Ash2 histone H3-K4 methyltransferase are tethered together selectively by the cell-proliferation factor HCF-1. Genes Dev 17: 896–911.

Wysocka J, Swigut T, Xiao H, Milne TA, Kwon SY, Landry J, Kauer M, Tackett AJ, Chait BT, Badenhorst P et al. 2006. A PHD finger of NURF couples histone H3 lysine 4 trimethylation with chromatin remodelling. Nature 442: 86–90.

Yang X, Qian K. 2017. Protein O-GlcNAcylation: emerging mechanisms and functions. Nat Rev Mol Cell Biol 18: 452–465.

Ying QL, Wray J, Nichols J, Batlle-Morera L, Doble B, Woodgett J, Cohen P, Smith A. 2008. The ground state of embryonic stem cell self-renewal. Nature 453: 519–523.

Zhang H, Chen Z, Ye Y, Ye Z, Cao D, Xiong Y, Srivastava M, Feng X, Tang M, Wang C et al. 2019. SLX4IP acts with SLX4 and XPF-ERCC1 to promote interstrand crosslink repair. Nucleic Acids Res 47: 10181–10201.

