## Supplemental Figures for "The BOD1L subunit of the SETD1A complex sustains the expression of DNA damage repair genes despite restraining H3K4 trimethylation"

### Materials and Methods

*Additional on targeting constructs, gene targeting and BAC transgenes*

The engrailed-intron-splice-acceptor-IRES-LacZ-Neomycin-polyA cassette flanked by FRT sites for FLP-recombination (Testa et al. 2004) was inserted in *Bod1l* intron 1. The critical exon 2, which upon deletion results in a frameshift and a premature stop codon in exon 3, was flanked by loxP sites. For targeting the second allele of *Bod1l*, the FRT flanked cassette was first removed from the targeted allele by transient expression of FLP from CAG-FLPo-IRES-puro (Kranz et al. 2010), then the *Bod1l* targeting construct was altered by recombineering to replace the neomycin resistance gene with blasticidin. The *Bod1l* targeting construct homology arms were 5 kb 5’ and 4.9 kb 3’.

Using recombineering, the BACs were first modified by exchanging the loxP site embedded in the BAC vector with the G418/kanamycin resistance gene expressed from the dual eukaryotic/prokaryotic SV40/Tn5 promoters (Fu et al. 2010) to enable G418 selection for the BACs in ESCs. *Bod1l, Bod1, Cxxc1* and *Bptf* were then tagged with Venus at their C-terminae using T2A/Gb3/blasticidin. The N-terminally Venus tagged *Setd1a* and *Setd1b* BAC transgenes (Bledau et al. 2014) and N-terminally tagged Mll1, Mll2 (Denissov et al. 2014) have been described. The X1, X3 and X4 deletions in the Venus-tagged *Setd1a* BAC were accomplished by recombineering to delete chosen exons by replacement with an ampicillin resistance gene to ensure that the reading frame is retained. The intron positions used for these deletions are indicated on Supplemental SFig2.

*Cell Proliferation and Cell Cycle Analysis*

The proliferation count assay was performed as described (Lubitz et al. 2007). For cell cycle analysis, asynchronously growing ES cells were fixed with ethanol and stained with 1 µg/ml propidium iodide (PI) containing 0.1 mg/ml RNase A (both from Sigma-Aldrich). Cells were analyzed by flow cytometry to determine G1, S, and G2/M cell cycle distribution with ModFit LT™ software (Verity Software House, Topsham, ME).

*esiRNA*

Endoribonuclease-prepared short interfering RNA (esiRNA) was synthesized according to the described protocol (Kittler et al. 2004). For transfection, ES cells were plated in gelatin-coated 24-well dishes and grown in complete medium without antibiotics to achieve 60-70% confluency on day 1. Then, 150 ng of esiRNA, which was diluted in 50 µl of serum-free DMEM, was mixed with 1 µl of Lipofectamine 2000 diluted in 50 µl of serum-free DMEM. After incubation for 20 minutes at room temperature, the mixture was added to freshly trypsinized ES cell culture suspended in 400 µl of complete medium without antibiotics but with serum and LIF. The medium was changed on day 2 and cells were harvested for Western Blot analysis 72 hours post-transfection.

*Computational methods*

AlphaFold (Jumper et al. 2021) was used to predict the X3 region alone (Figure 1). AlphaFold-Multimer (Abramson et al. 2024) was used to predict the co-folded interactions of BOD1L-SETD1A, BOD1-SETD1B (Supplemental Fig S3) and BOD1-PP2A-5B (interacting with residues 433 to 477, Supplemental Fig S4). Pymol v.2.3.0 was utilised for visualisation and structural alignment using the CEalign method. Non-covalent interactions were identified using PLIP (Protein-Ligand Interaction Profiler; (Adasme et al. 2021). Sequence identity was determined using the BLAST (Basic Local Alignment Search Tool) platform (as of November 28, 2022). Both BOD1L-SETD1A, BOD1-SETD1B complexes achieved a very high confidence score of 80 to 90%. In a second step, we structurally aligned both complexes (Supplemental Fig S3D). The alignment is very comprehensive with 176 out of 200 residues aligned. It has an overall root-mean-square deviation (RMSD) of 1.4Å and a template modelling (TM ) score of 0.65, where values greater than 0.5 indicate a high likelihood of common ancestry (Xu and Zhang 2010). However, the BOD1-PP2A-5B confidence score was only 40 to 50% for the helix, which meant that the template modelling approach for further validation was not viable.

The multiple sequence alignment of SET1 family members (Supplemental Fig. S2B) was obtained with MAFFT (Katoh et al. 2005) with default parameters as implemented in Jalview (Waterhouse et al. 2009) and colour coded for conservation using the Clustal X colour scheme (Gibson et al. 1994). The sequences were selected to emphasize both conservation and diversity in the SET1 protein family.

### Supplemental References

Abramson J, Adler J, Dunger J, Evans R, Green T, Pritzel A, Ronneberger O, Willmore L, Ballard AJ, Bambrick J et al. 2024. Accurate structure prediction of biomolecular interactions with AlphaFold 3. *Nature* **630**: 493-500.

Adasme MF, Linnemann KL, Bolz SN, Kaiser F, Salentin S, Haupt VJ, Schroeder M. 2021. PLIP 2021: expanding the scope of the protein-ligand interaction profiler to DNA and RNA. *Nucleic Acids Res* **49**: W530-W534.

Bledau AS, Schmidt K, Neumann K, Hill U, Ciotta G, Gupta A, Torres DC, Fu J, Kranz A, Stewart AF et al. 2014. The H3K4 methyltransferase Setd1a is first required at the epiblast stage, whereas Setd1b becomes essential after gastrulation. *Development* **141**: 1022-1035.

Denissov S, Hofemeister H, Marks H, Kranz A, Ciotta G, Singh S, Anastassiadis K, Stunnenberg HG, Stewart AF. 2014. Mll2 is required for H3K4 trimethylation on bivalent promoters in embryonic stem cells, whereas Mll1 is redundant. *Development* **141**: 526-537.

Fu J, Teucher M, Anastassiadis K, Skarnes W, Stewart AF. 2010. A recombineering pipeline to make conditional targeting constructs. *Methods Enzymol* **477**: 125-144.

Gibson TJ, Hyvonen M, Musacchio A, Saraste M, Birney E. 1994. PH domain: the first anniversary. *Trends Biochem Sci* **19**: 349-353.

Hoshii T, Cifani P, Feng Z, Huang CH, Koche R, Chen CW, Delaney CD, Lowe SW, Kentsis A, Armstrong SA. 2018. A Non-catalytic Function of SETD1A Regulates Cyclin K and the DNA Damage Response. *Cell* **172**: 1007-1021 e1017.

**26**: 889-895.

Jumper J, Evans R, Pritzel A, Green T, Figurnov M, Ronneberger O, Tunyasuvunakool K, Bates R, Zidek A, Potapenko A et al. 2021. Highly accurate protein structure prediction with AlphaFold. *Nature* **596**: 583-589.
Katoh K, Kuma K, Toh H, Miyata T. 2005. MAFFT version 5: improvement in accuracy of multiple sequence alignment. *Nucleic Acids Res* **33**: 511-518.
Kittler R, Putz G, Pelletier L, Poser I, Heninger AK, Drechsel D, Fischer S, Konstantinova I, Habermann B, Grabner H et al. 2004. An endoribonuclease-prepared siRNA screen in human cells identifies genes essential for cell division. *Nature* **432**: 1036-1040.
Kranz A, Fu J, Duerschke K, Weidlich S, Naumann R, Stewart AF, Anastassiadis K. 2010. An improved Flp deleter mouse in C57Bl/6 based on Flpo recombinase. *Genesis* **48**: 512-520.
Lubitz S, Glaser S, Schaft J, Stewart AF, Anastassiadis K. 2007. Increased apoptosis and skewed differentiation in mouse embryonic stem cells lacking the histone methyltransferase Mll2. *Mol Biol Cell* **18**: 2356-2366.
Testa G, Schaft J, van der Hoeven F, Glaser S, Anastassiadis K, Zhang Y, Hermann T, Stremmel W, Stewart AF. 2004. A reliable lacZ expression reporter cassette for multipurpose, knockout-first alleles. *Genesis* **38**: 151-158.
Waterhouse AM, Procter JB, Martin DM, Clamp M, Barton GJ. 2009. Jalview Version 2--a multiple sequence alignment editor and analysis workbench. *Bioinformatics* **25**: 1189-1191.
Xu J, Zhang Y. 2010. How significant is a protein structure similarity with TM-score = 0.5? *Bioinformatics*

### Supplemental Figures

#### Suppl Fig S1

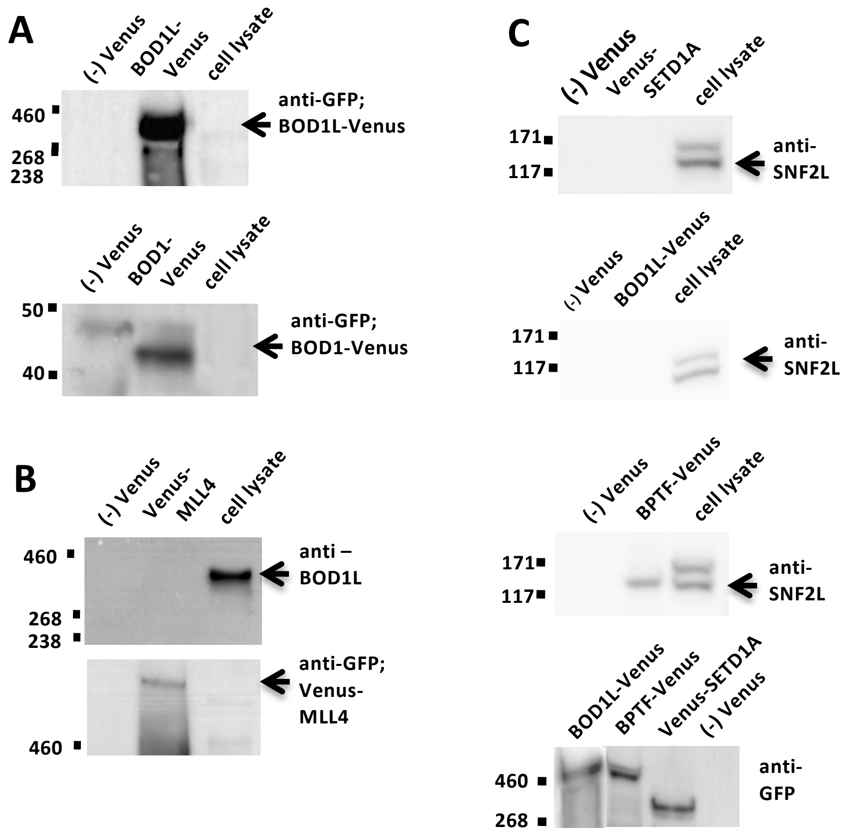


**Supplemental Figure S1. Co-immunoprecipitation of BOD1L-Venus, BOD1-Venus, BPTF-Venus, Venus-SETD1A, Venus-MLL4 protein lysates.**

**A.** Immunoprecipitation controls for BOD1L-Venus and BOD1-Venus.

**B.** Immunoprecipitation of Venus-MLL4 probed with either BOD1L or anti-GFP antibodies.

**C.** Immunoprecipitation of Venus-SETD1A, BOD1L-Venus and BPTF-Venus probed with anti-SNF2L antibody. At bottom, immunoprecipitation control probed with anti-GFP antibody.

SETD1A, observed ≈ 250 kDa; Venus-SETD1A ≈ 280 kDa; BOD1L-Venus ≈ 360 kDa; BPTF ≈ 330 kDa; BPTF-Venus ≈ 360 kDa; SNF2L≈ 120 kDa.

**Suppl Fig 2**

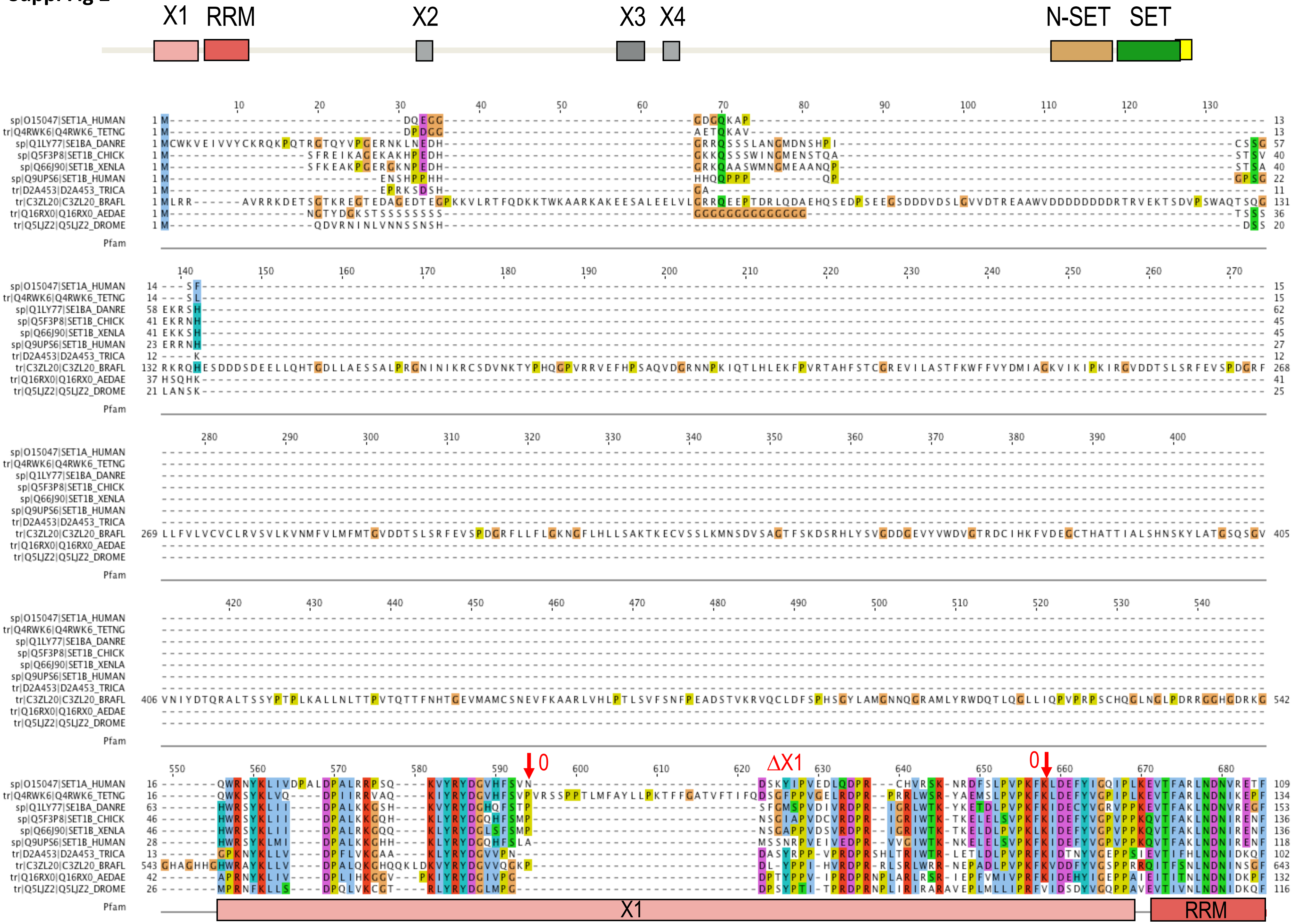
X1
RRM
X2
X3
X4
N-SET
SET
sp|O15047|SET1A_HUMAN
tr|Q4RWK6|Q4RWK6_TETNG
sp|Q1LY77|SE1BA_DANRE
sp|Q5F3P8|SET1B_CHICK
sp|Q66J90|SET1B_XENLA
sp|Q9UPS6|SET1B_HUMAN
tr|D2A453|D2A453_TRICA
tr|C3ZL20|C3ZL20_BRAFL
tr|Q16RX0|Q16RX0_AEDAE
tr|Q5LJZ2|Q5LJZ2_DROME
Pfam
0
ΔX1
0
X1
RRM

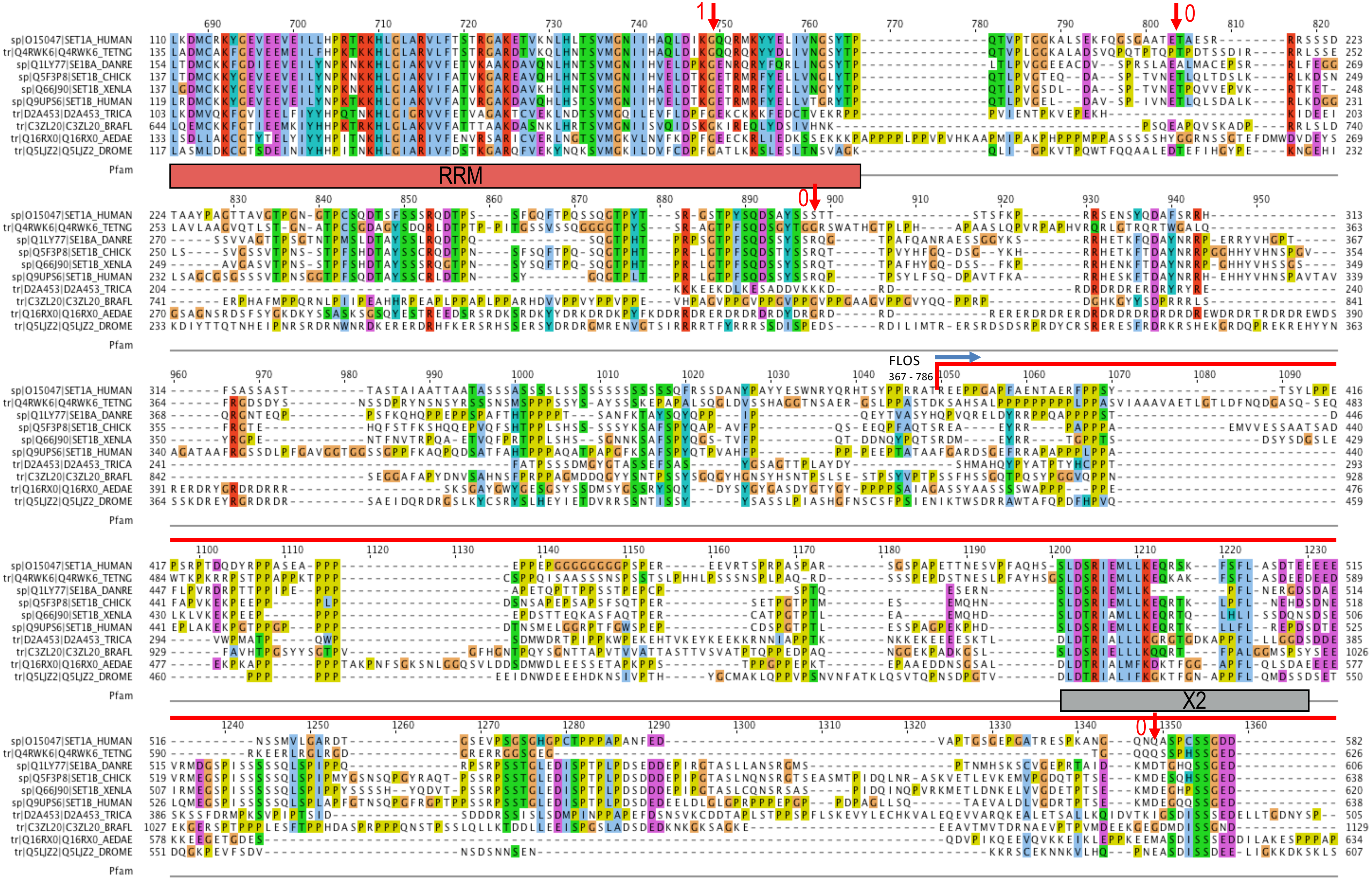
sp|O15047|SET1A_HUMAN
tr|Q4RWK6|Q4RWK6_TETNG
sp|Q1LY77|SE1BA_DANRE
sp|Q5F3P8|SET1B_CHICK
sp|Q66J90|SET1B_XENLA
sp|Q9UPS6|SET1B_HUMAN
tr|D2A453|D2A453_TRICA
tr|C3ZL20|C3ZL20_BRAFL
tr|Q16RX0|Q16RX0_AEDAE
tr|Q5LJZ2|Q5LJZ2_DROME
Pfam
RRM
FLOS
367 - 786
X2
1
0

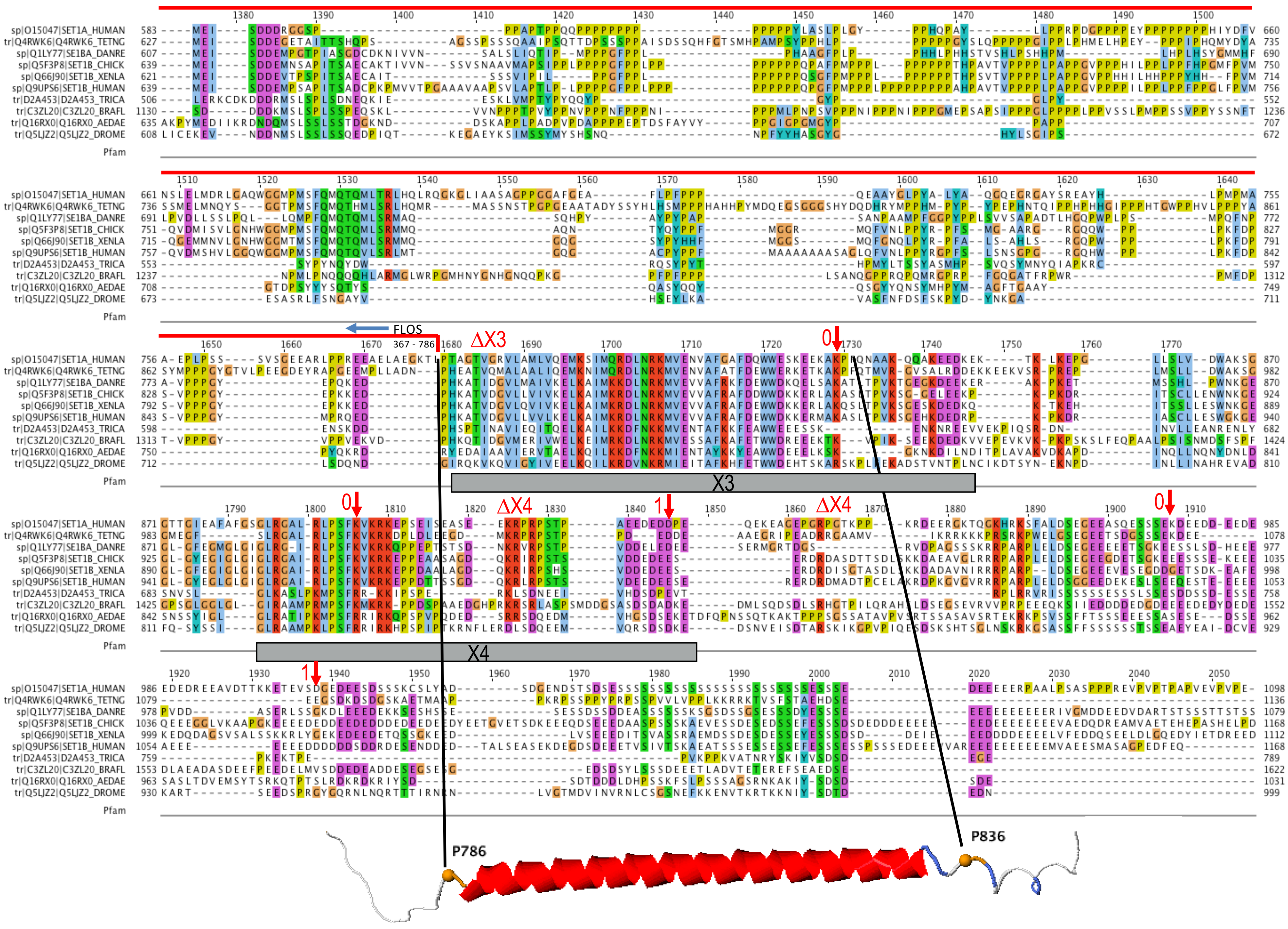

sp|O15047|SET1A_HUMAN
tr|Q4RWK6|Q4RWK6_TETNG
sp|Q1LY77|SE1BA_DANRE
sp|Q5F3P8|SET1B_CHICK
sp|Q66J90|SET1B_XENLA
sp|Q9UPS6|SET1B_HUMAN
tr|D2A453|D2A453_TRICA
tr|C3ZL20|C3ZL20_BRAFL
tr|Q16RX0|Q16RX0_AEDAE
tr|Q5LJZ2|Q5LJZ2_DROME
Pfam
FLOS
367 - 786
ΔX3
ΔX4
X3
X4
0
1
P786
P836

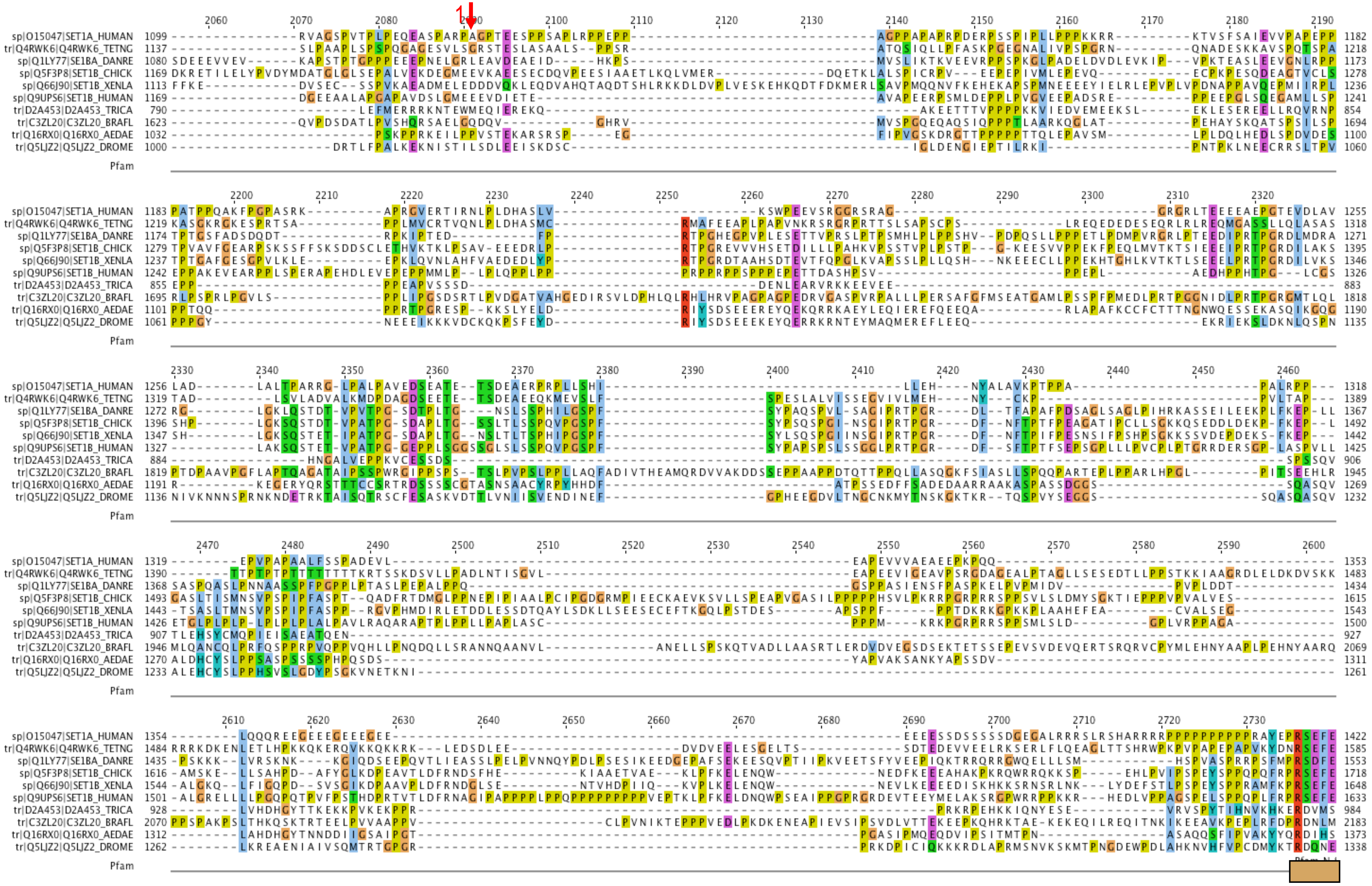
sp|O15047|SET1A_HUMAN
tr|Q4RWK6|Q4RWK6_TETNG
sp|Q1LY77|SET1B_DANRE
sp|Q5F3P8|SET1B_CHICK
sp|Q66J90|SET1B_XENLA
sp|Q9UPS6|SET1B_HUMAN
tr|D2A453|D2A453_TRICA
tr|C3ZL20|C3ZL20_BRAFL
tr|Q16RX0|Q16RX0_AEDAE
tr|Q5LJZ2|Q5LJZ2_DROME
Pfam

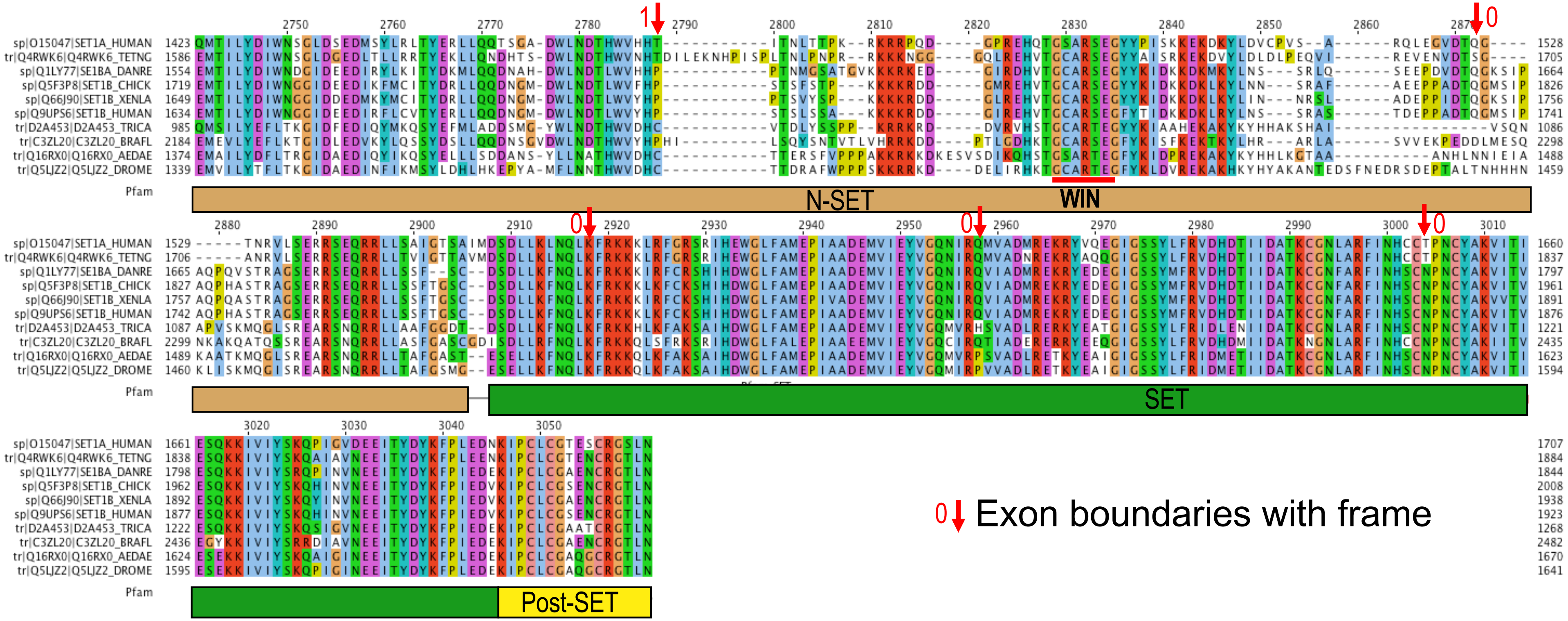


**Supplemental. Fig. S2. Multiple sequence alignment of the SET1 family.**

Multiple sequence alignment of selected members of the SET1 family, including human SETD1A and B (SET1A_HUMAN; SET1B_HUMAN); zebrafish SETD1B-A (SE1BA_DANRE); chicken SETD1B (SET1B_CHICK); frog SETD1B (SET1B_XENLA); fruit fly SET1 (Q5LJZ2_DROME); and SET1 family members from pufferfish (Q4RWK6_TETNG), tribolium (D2A453_TRICA), lancelet (C3ZL20_BRAFL), and yellow fever mosquito (Q16RX0_AEDAE). UniProt IDs are given in parentheses. A generic cartoon of the protein family is depicted at the top displaying known protein domains (RRM, N-SET and SET) as well as four additional, conserved regions termed X1-X4. These domains and regions are also indicated underneath the alignment. The AlphaFold prediction for X3 is also displayed. The region called FLOS (residues 389-785 in SET1A_HUMAN; Hoshii et al., 2018) is marked with a red line and arrows above the alignment. Conserved exon boundaries are indicated with vertical red arrows and the reading frame. WIN – WDR5 interaction site.

**Suppl Fig S3**

**A** BOD1-SETD1B

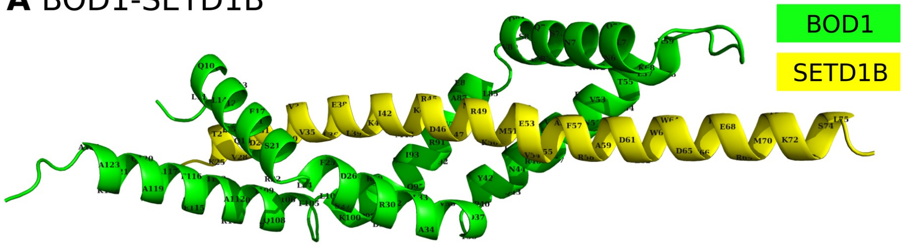


**B** BOD1L-SETD1A

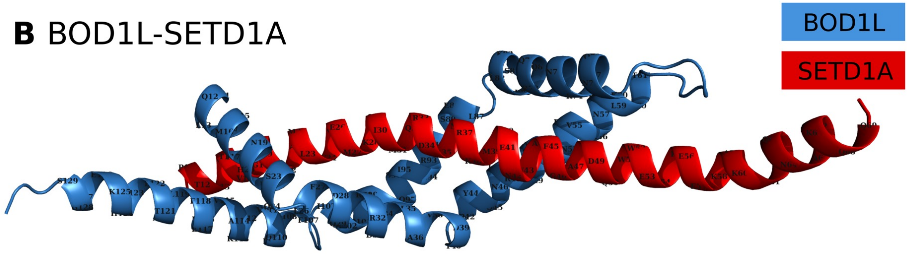


**C** BOD1-SETD1B

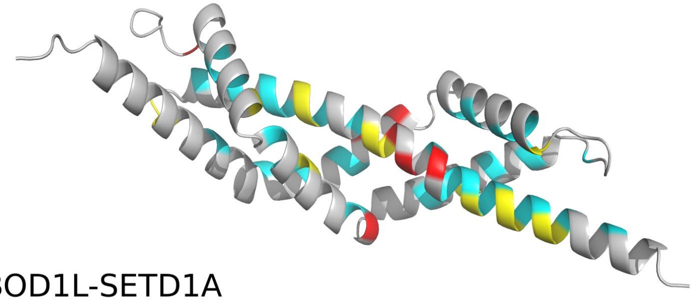

**D** BOD1L-SETD1A

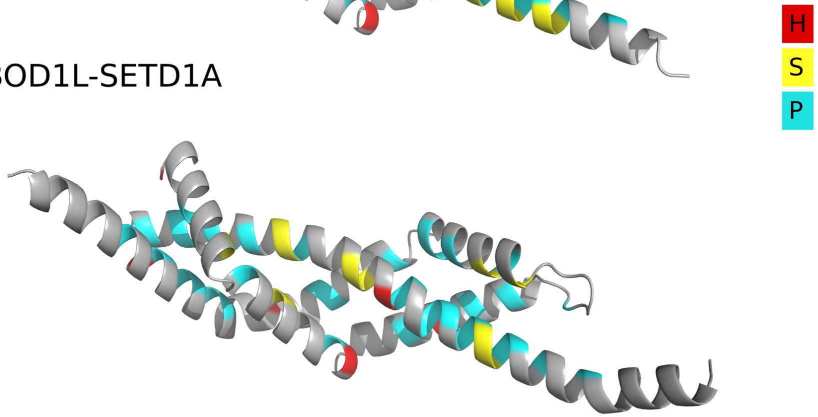

**Suppl Fig S3**

**E**

```
BOD1L
          20        30        40        50        60        70        80        90       100       110       120
DPQLVAMIVNHLKSQGLFDQFRRDCLADVDTKPAYQNLRQRVDNFVANHLATHTWSPHLNKNQLRNNIRQQVLKSGMLESGIDRIISQVVDPKINHTFRPQVEKAVHEFLATLNHKEEGS
-------P----S----P---S------PH--------P---H--P---H----P-----S--PS---P--PP----P------PP----H------PP--PP--PH--P----------
****.*.**  ***.**** ********************.*****. **    *.* .******* .** *..*****.*.***********.** ****.*.*.*****
----P--PP---S----P--PS------PH--------P---S--P---H----P-----S--PP---H--P------H---H-PP---P---P-------P----S-------------
DPQLIALIVEQLKSRGLFDSFRRDCLADVDTKPAYQNLRQKVDNFVSTHLDKQEWNPTMNKNQLRNGLRQSVVQSGMLEAGVDRIISQVVDPKLNHIFRPQIERAIHEFLAAQKKAAVPA
  10        20        30        40        50        60        70        80        90       100       110       120
BOD1


SETD1A
         10        20        30        40        50        60
EAELAEGKLPTAGTVGRVLAMLVQEMKSIMQRDLNRKMVENVAFGAFDQWWESKEEKAKPFQNA
----------H--PP-HPP--P--SSH-----SS--HPP--HPP-PPSSPP--P---P------
.      . *   **  **  ....*.*.**.********* *** ***.**. **  **
--------SS---H---P--PPP-SSP---H-SSH-H-PH-PPPSPPSSHPSSP---P------
QDTSPRQEDPHKATVDGVLLVVLKELKAIMKRDLNRKMVEVVAFRAFDEWWDKKERMAKASLTP
      20           30        40        50        60        70
SETD1B
```

**Supplemental Fig. S3. AlphaFold predictions of BOD1L/SETD1A-X3 and BOD1/SETD1B-X3 interactions.**

**A.** The Shg1 box from BOD1 was fused to the X3 helix from SETD1B and subjected to AlphaFold analysis. The linker sequence in between has been omitted for clarity. Green BOD1; Yellow SETD1B-X3. Selected residues are labelled.

**B.** Same as A except for BOD1L and SETD1A. Blue BOD1L; Red SETD1A-X3.

**C.** Binding interactions for BOD1-SETD1B-X3; red-hydrogen bonds; yellow-salt bridges; cyan-hydrophobic contact.

**D.** Same as C except for BOD1L-SETD1A-X3.

**E**. Sequence alignments of BOD1L and BOD1 (above) and SETD1A-X3 and SETD1B-X3 (below) showing (i) in the center, a star for identity and a dot for a positive Blosum score, and (ii) denoted interactions; H for hydrogen bond, P for hydrophobic contact, S for salt bridge.

**A** PP2A-5B (56kD beta, Q15173)

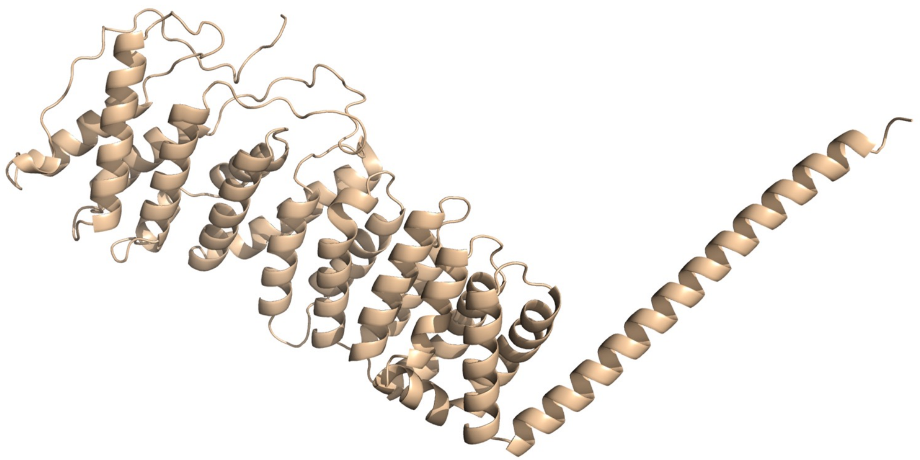

**B** PP2A-5B (56kD beta, Q15173)

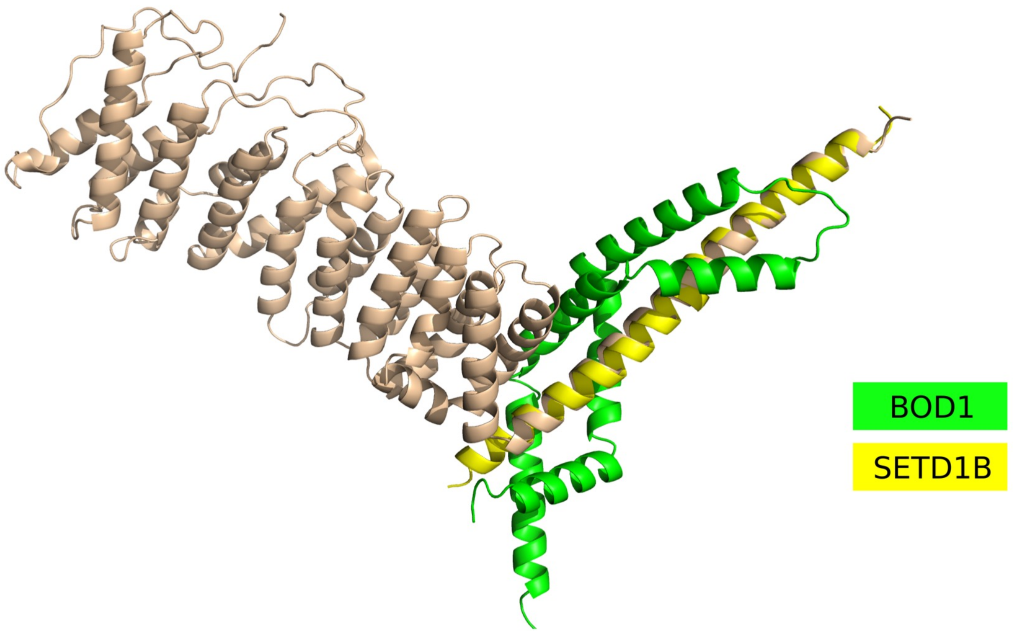


**Supplemental Fig. S4.** AlphaFold prediction of the BOD1 – PP2A-5B interaction.

**A.** AlphaFold prediction of PP2A-5B. The long PP2A-5B helix is C-terminal including a.a.s 433 - 477.

**B.** The predicted BOD1 binding provokes a clash with PP2A-5B sequences N-terminal to the helix, suggesting that a conformational change is required for binding. BOD1 – green; SETD1B – yellow imposed over the PP2A-5B helix.

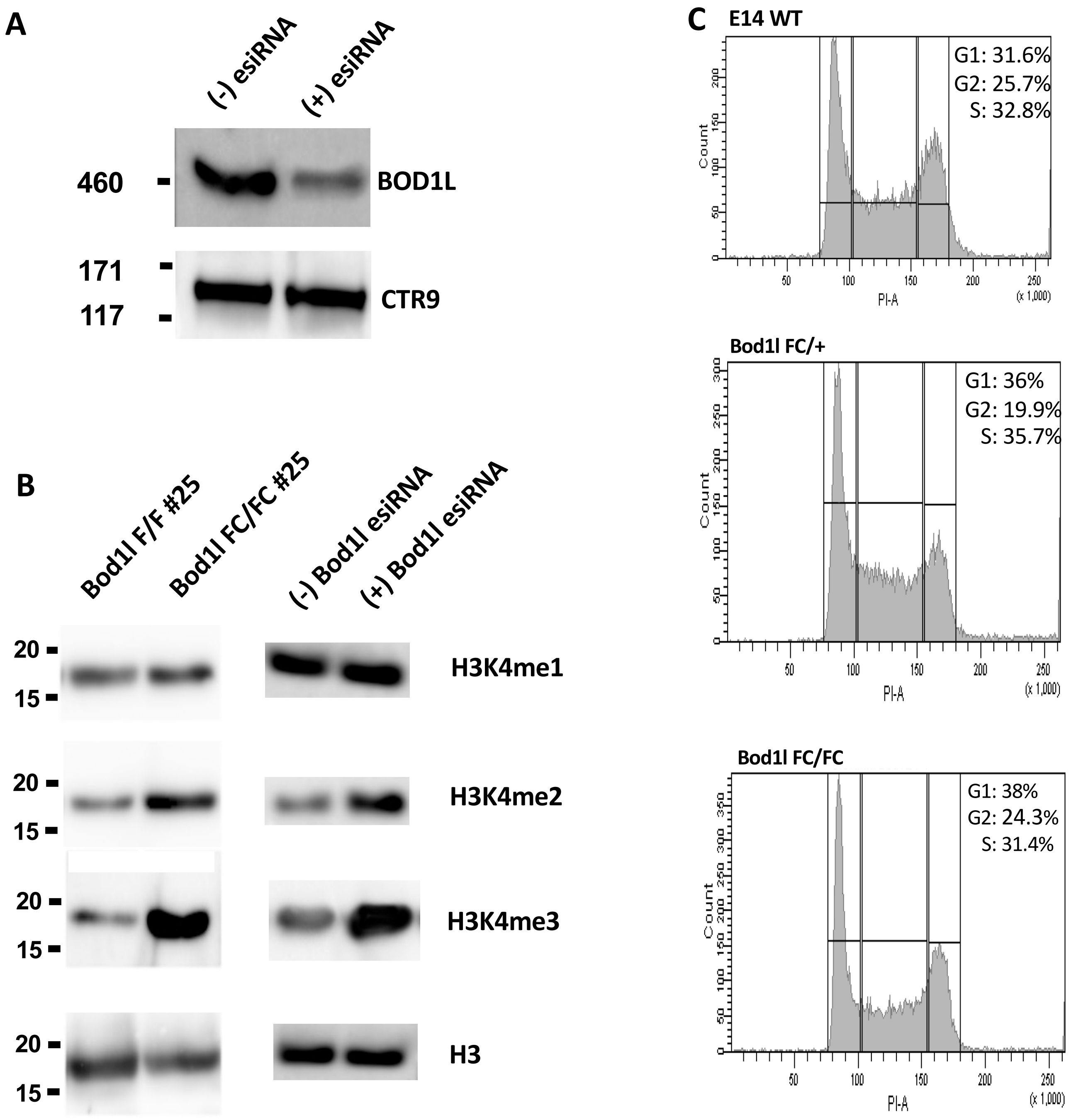


**Supplemental Fig. S5. RNAi knockdown of Bod1l in E14 ESCs**

**A.** Western analysis of extract harvested 72 hours after administration of esiRNA against BOD1L using anti-BOD1L antibody and anti-CTR9 antibody as control for input.

**B.** Western analysis of histone 3 96 hours after adding 4-OH tamoxifen or not (left columns) or 72 hours after adding Bod1l esiRNA or not (right columns).

**C.** FACS analysis of propidium iodide stained E14, Bod1l FC/+ and Bod1l FC/FC ESCs.

SFig 6A

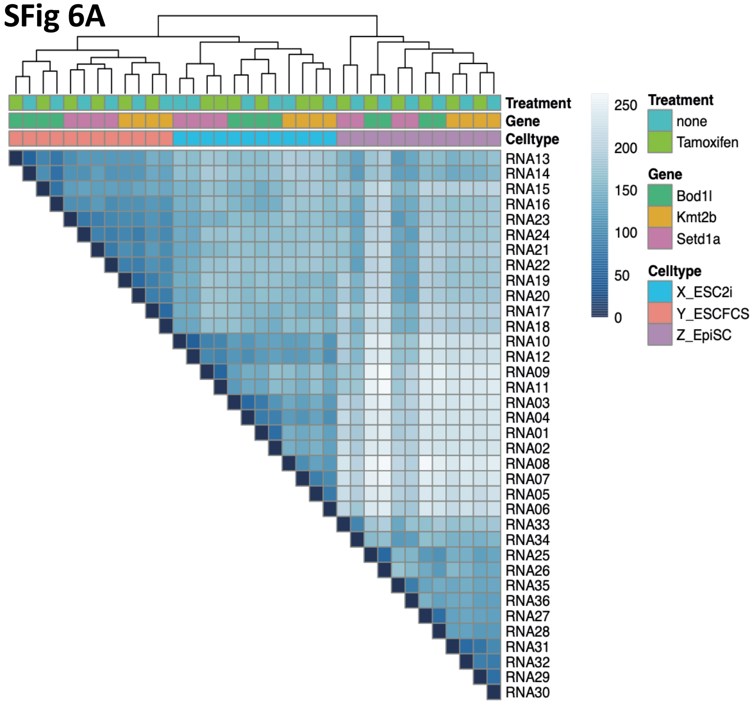


SFig 6B

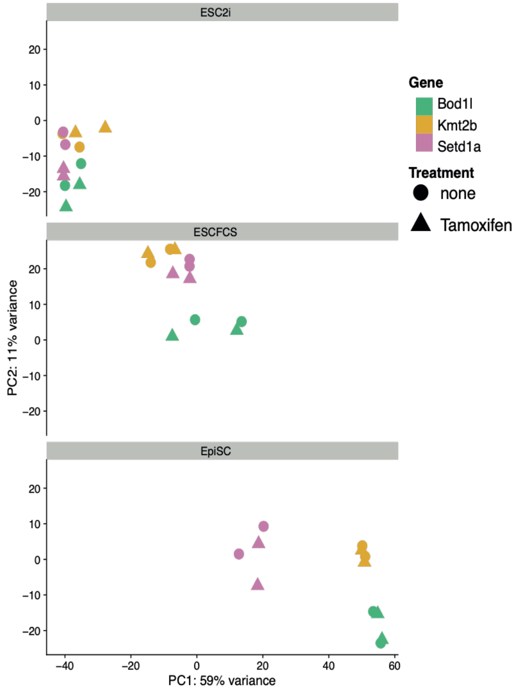


**Supplemental Fig. S6. Further analysis of RNA-seq**

**A.** Hierarchical clustering of the 36 RNA-seq datasets indicates a high degree of technical quality.

**B.** Principal component analysis of the 36 RNA-seq datasets. Duplicates of *Bod1l* F/F, *Setd1a* F/F and *Mll2/Kmt2b* F/F were harvested for mRNA 48 hours after treating for 48 hours with 4 hydroxy tamoxifen or not (total 96 hours) from ESCs grown either in the presence of the two inhibitors and LIF (2i), in the presence of FCS and LIF (FCS), or in the absence of LIF but presence of FGF2 and Activin A (Epi). All three duplicates were grown in parallel for each growth condition. Circles, no 4-hydroxy tamoxifen; triangles, with 4-hydroxy tamoxifen.

**C.** Pairwise comparisons of gene expression from the three cellular conditions using conditional knock-out dependent log2 fold changes from the differential analysis. Genes are highlighted when significant with an adjusted p-value $< 0.05$ within the stated condition.

**D.** Heatmap hierarchical clustering based on genes that respond equivalently to loss of *Setd1a* or *Bod1l*. Left panel – genes differentially deregulated dependent on both Setd1a and Bod1l; right panel – genes specifically differentially regulated dependent on Setd1a, but not Bod1l.

**E**. H3K4me3 peaks associated with the 12,996 active TSSs in ESCFCSs in wild type (blue) and Bod1 FC/FC after treating for 48 hours with 4 hydroxy tamoxifen or not (total 96 hours). Below are subsets involving the most strongly downregulated DEGs (144), the genes listed in Table 1 (51) and the genes that occur in Table 1in at least two columns after Bod1l knockout (27).

**Suppl Fig S6C**

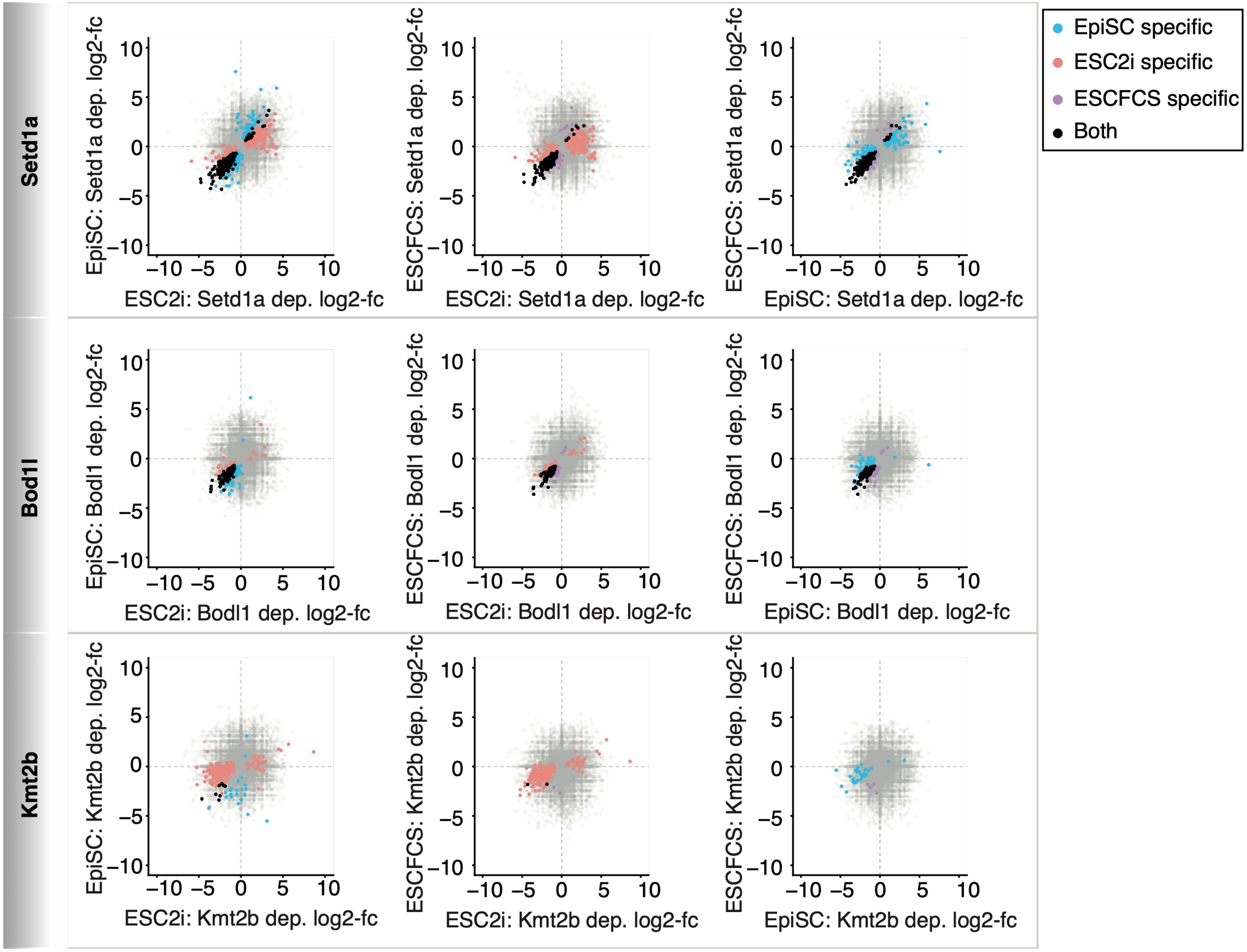

D

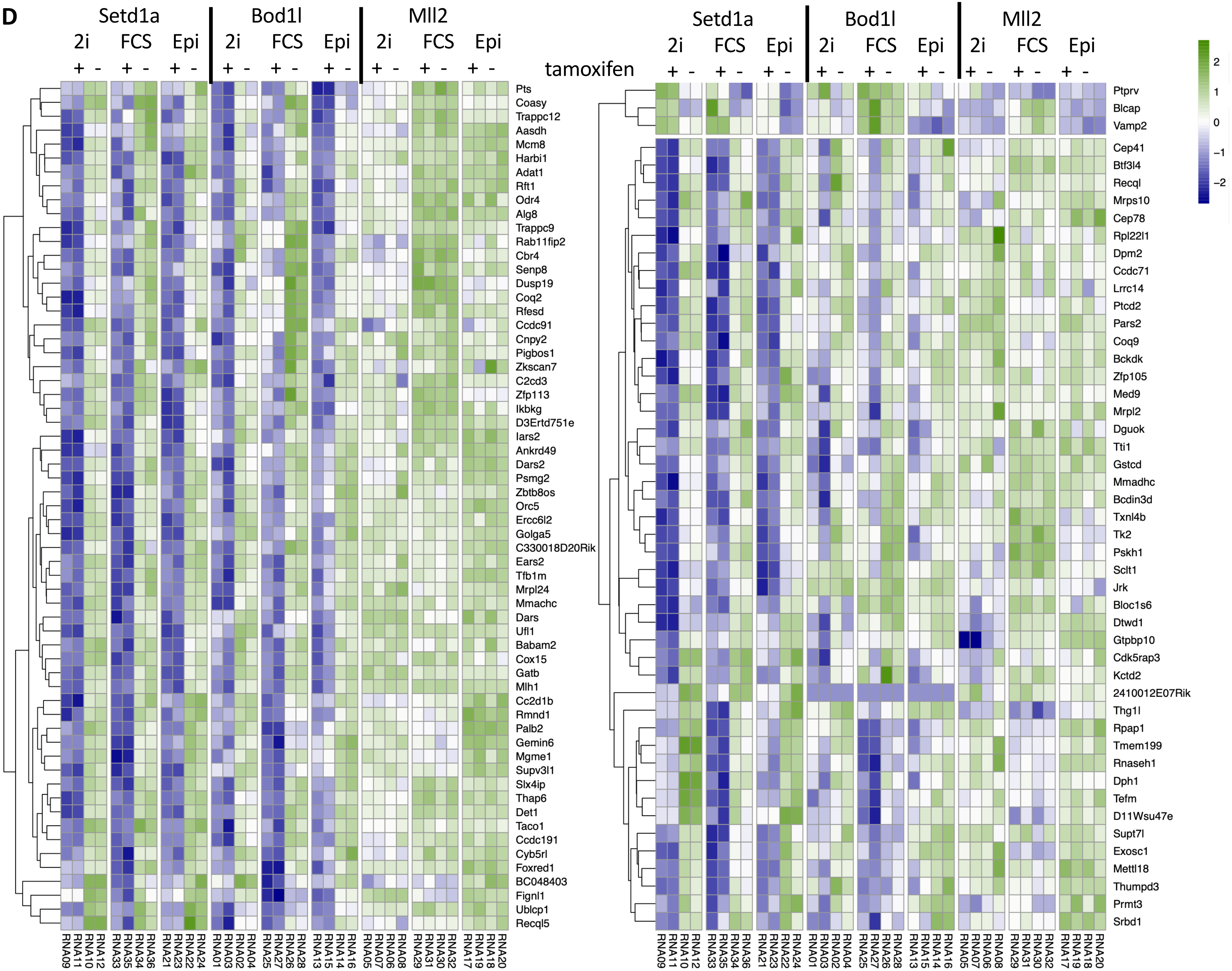
Setd1a
Bod1l
Mll2
2i
FCS
Epi
+ -
tamoxifen
Pts
Coasy
Trappc12
Aasdh
Mcm8
Harbi1
Adat1
Rft1
Odr4
Alg8
Trappc9
Rab11fip2
Cbr4
Senp8
Dusp19
Coq2
Rfesd
Ccdc91
Cnpy2
Pigbos1
Zkscan7
C2cd3
Zfp113
Ikbkg
D3Ertd751e
Iars2
Ankrd49
Dars2
Psmg2
Zbtb8os
Orc5
Ercc6l2
Golga5
C330018D20Rik
Ears2
Tfb1m
Mrpl24
Mmachc
Dars
Ufl1
Babam2
Cox15
Gatb
Mlh1
Cc2d1b
Rmnd1
Palb2
Gemin6
Mgme1
Supv3l1
Slx4ip
Thap6
Det1
Taco1
Ccdc191
Cyb5rl
Foxred1
BC048403
Fignl1
Ublcp1
Recql5
Ptprv
Blcap
Vamp2
Cep41
Btf3l4
Recql
Mrps10
Cep78
Rpl22l1
Dpm2
Ccdc71
Lrrc14
Ptcd2
Pars2
Coq9
Bckdk
Zfp105
Med9
Mrpl2
Dguok
Tti1
Gstcd
Mmadhc
Bcdin3d
Txnl4b
Tk2
Pskh1
Sclt1
Jrk
Bloc1s6
Dtwd1
Gtpbp10
Cdk5rap3
Kctd2
2410012E07Rik
Thg1l
Rpap1
Tmem199
Rnaseh1
Dph1
Tefm
D11Wsu47e
Supt7l
Exosc1
Mettl18
Thumpd3
Prmt3
Srbd1
2
1
0
−1
−2
RNA09
RNA11
RNA10
RNA12
RNA33
RNA35
RNA34
RNA36
RNA21
RNA23
RNA22
RNA24
RNA01
RNA03
RNA02
RNA04
RNA25
RNA27
RNA26
RNA28
RNA13
RNA15
RNA14
RNA16
RNA05
RNA07
RNA06
RNA08
RNA29
RNA31
RNA30
RNA32
RNA17
RNA19
RNA18
RNA20

**E**

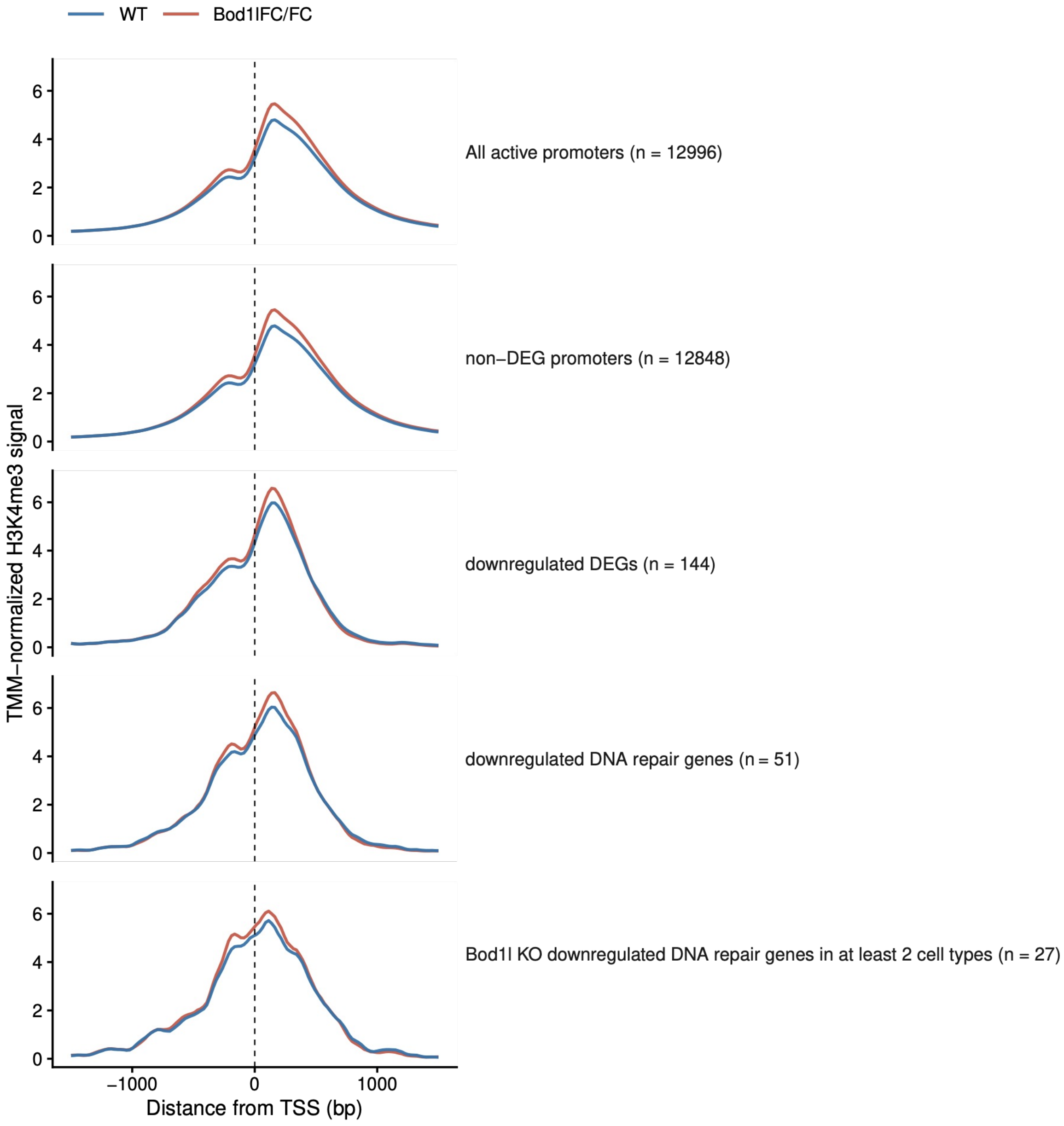

WT
Bod1lFC/FC
TMM−normalized H3K4me3 signal
All active promoters (n = 12996)
non−DEG promoters (n = 12848)
downregulated DEGs (n = 144)
downregulated DNA repair genes (n = 51)
Bod1l KO downregulated DNA repair genes in at least 2 cell types (n = 27)
−1000
0
1000
Distance from TSS (bp)
