## Supplementary material for "The BOD1L subunit of the SETD1A complex sustains the expression of DNA damage repair genes despite restraining H3K4 trimethylation": Response to Reviewers

### Full Revision

Review COMMONS

**Manuscript number:** RC-2026-03448
**Corresponding author(s):** A. Francis Stewart

#### 1. General Statements [optional]

On behalf of the authors, I thank the reviewers for their critical reading of the manuscript. We really appreciate the care and attention they have applied to their reading and reports.
An initial comment may add some context - in accordance with German law, I (AFS) had to retire from my position at Dresden University in October 2023. I am still working however have very little capacity to add new experiments to the manuscript. Consequently my response to the reviewers is somewhat more critical than the normal concessions and acquiescence that are usually adopted.

#### 2. Point-by-point description of the revisions

**Reviewer #1 Summary**
**This manuscript describes functional characterization of Bod1 family proteins (particularly Bod1L) and their interaction with COMPASS family histone methyltransferase complexes. Bod1 proteins are conserved through evolution and related to yeast Shg1, which binds to the yeast COMPASS via a conserved domain and negatively regulates H3K4me3 levels. Here the authors performed IP-MS of Bod1L, Bod1, Setd1a, and Setd1b in engineered mouse embryonic stem cells and confirmed presence of Bod1 and Bod1L in both Setd1a and Setd1b-containing COMPASS complexes. AlphaFold modeling predicted an interaction between the Bod1/L Shg domain and a conserved helix in Setd1a/b which was validated in stable Setd1a deletion ESC lines and with isolated Sed1a/b fragments. Functional analysis of Bod1L and Setd1a deletion lines confirmed that Bod1L negatively regulated H3K4me3 levels. Moreover, the knockouts caused parallel effects on the expression of DNA repair genes and both apparently enhanced levels of baseline DNA damage, in agreement with findings in leukemia cell lines. This argues that Setd1a/Bod1L regulates DNA repair gene expression independently of H3K4me3.**

**Major comments**
**Figures 2D, 2E, 3C: there are no panels showing the efficiency of Bod1 or Bod1L immunoprecipitation. The immunoblots seem to indicate that either Bod1 or Bod1L precipitate a substantial fraction of Setd1a and a much smaller fraction of Setd1b, although it is impossible to tell without the blots of Bod1/Bod1L. The idea that Setd1a is primarily associated with Bod1L vs Bod1 is presumed in the rest of the manuscript (likely based on previous results in other cell lines) but is not strongly supported by these figures.**

Response: The protein complex and interaction data is based on AP-MS (affinity purification-mass spectrometry) and the results are presented as Volcano plots, which is the standard and

most accessible format. AP-MS acquires candidate data because some bona-fide interactions will be missed and spurious interactions will be included, especially when the threshold of significance is lowered. To validate the candidate data from tagged SETD1A and SETD1B we used
(a) reciprocal AP-MS with tagged BOD1L, BOD1 and CXXC1. These results secured the primary conclusion regarding the associations of BOD1L and BOD1 with SETD1A and SETD1B (as well as other subunits). I should add that we show – for the first time – functional evidence that BOD1L is a subunit of the SETD1A complex (Figure 5).
(b) immunoprecipitations to confirm selected interactions. As stated in the legend of Figure 2, 10% total extract are shown as controls. This control allows the reader to evaluate the efficiency of the associated protein in the IP. For example, (Fig. 2D), clearly BOD1L associates much more with SETD1A than with SETD1B. Nevertheless, the association with SETD1B was detected and reciprocally confirmed (Fig. 3C). Another example, (Fig. 2E), clearly BPTF interacts with BOD1L at notable efficiency but not with SETD1A. There are also additional IPs in the supplement that add further confidence.
Additionally, AP-MS with three SETD1A deletion mutants (Fig. 4) adds supporting evidence to the conclusions drawn from Figures 2 and 3. The implication that BOD1L and BOD1 interact with X3, which in Figure 4 is a negative result and therefore – because negative results in AP-MS cannot be used to draw conclusions – we explicitly tested the proposition that BOD1L and BOD1 interact with X3 to secure the conclusion (Fig 4D).
Consequently, with respect, we do not agree with the following comment by reviewer 1 -
**- although it is impossible to tell without the blots of Bod1/Bod1L. The idea that Setd1a is primarily associated with Bod1L vs Bod1 is presumed in the rest of the manuscript (likely based on previous results in other cell lines) but is not strongly supported by these figures.**

- because the data presented allows quantitative evaluation and consequently conclusions about the BOD1L and SET1A interactions. The primary association of BOD1L with SETD1A is established by the data presented, as is the secondary associations between BOD1 and SETD1A, as well as BOD1L and SETD1B.

**2. Figure 4: the SETD1A-X1 and X3 internal deletion lines show many interesting interactions not detected in the wild-type line, notably with CPSF components. What is the significance of this?**
AP-MS explores the proteome and a number of intriguing associations can be found in addition to the most robust biochemical interactions. In this manuscript, we present some exploration of the intriguing extras but remain focused on the SETD1A and B complexes. We and others have published on the connection between the yeast Set1C and yeast CPSF, but exploring that issue connection is beyond the experimental and conceptual themes of this manuscript, especially considering other comments by the reviewers about reducing the manuscript.

**3. Related to Figure 4: Why were none of the functional genomics experiments described in Figures 5-7 performed on the Setd1a-X3 deletion line given that the authors had it in hand? This would have been a logical complement to the Bod1L deletion experiment and further**

**addressed the issue of functional partnership between Setd1a and Bod1/L. One could make a similar point regarding Bod1-it is unclear why (given the co-IP and AP-MS data in Figures 2 and 3) a deletion line of Bod1 was not analyzed in parallel. There was convincing rationale in the Hoshii 2024 paper to focus on Setd1a/Bod1L in that system; the rationale for doing this here is less clear.**

We thank the reviewer for this suggestion. Indeed this is a good experiment that emerges from the data we are presenting (and not from Hoshii et al 2024). We take this excellent suggestion as an indication that our manuscript has presented the evidence sufficiently well to permit the reviewer to make this suggestion, which is worth pursing in a new project. However the manuscript is already replete with progress. It is worth mentioning that in response to an earlier round of reviewing elsewhere, we added the experiment that is now Figure 7. This process of adding further experiments in response to thoughtful reviewing comes at the risk of promoting further good suggestions, which are constructive and welcome but at some point progress should be published.

**4. Figure 5G: This figure does not seem to include a control in which wild-type ESCs are treated with tamoxifen in parallel with the Flp Bod1L line.**

There is no published or conceptual reason to include a control for the induction of DNA damage by tamoxifen in wild type cells. My lab pioneered ligand inducible conditional mutagenesis (Logie C and Stewart AF. 1995 Ligand-regulated site-specific recombination. PNAS 92, 5940-5944) including tamoxifen inducible conditional mutagenesis in mice (Schwenk F, Kühn R, Angrand P-O, Rajewsky K and Stewart AF. 1998 Temporally and spatially regulated somatic mutagenesis in mice. Nucleic Acids Res. 26, 1427-1432) and we have published advice for the appropriate controls for tamoxifen induced conditional mutagenesis (Anastassiadis et al 2010, Methods Enzymol, 477, 109-23). All experiments involving tamoxifen induction of conditional mutagenesis were thoroughly accompanied by appropriate controls. In the case of Fig. 5G, administration of tamoxifen to ESCs had no detectable effect on DNA damage as evaluated by p-H2AX or p-ATM staining – as expected and therefore not shown.

**5. Figure 6: This figure should include Venn diagrams that clearly show the overlap between genes affected by Bod1L removal compared to Setd1a.**

In Figure 6B, the overlap is clearly illustrated in a colour presentation that we think is superior to presenting these data as a Venn diagram. These data are also presented in different formats - in the Supplement Fig. 6D and the most significant DNA repair genes are listed in Table 1 again presented in an overlapping format.

**6. Related to Figure 6/7: These figures should include analysis of the H3K4me3 ChIP-seq data in Figure 5 specifically at Bod1L-regulated DEGs.**

We now present a new Supplemental Figure 6E and include the statement - ‘Increased H3K4me3 peaks were also observed at the promoters of the DNA repair genes that showed decreased expression after loss of BOD1L (Supplemental Figure 6E).’ in the text. In other words, elevated H3K4me3 is also observed on the DNA repair genes that show decreased expression.

**7. Discussion: The authors should note that the direct role of Setd1a at DNA breaks is proposed to rely on enzymatic activity (Higgs et al 2018, Bayley et al 2022).**
A sentence regarding SETD1A enzyme activity in DNA repair has been included in the Discussion.

**Minor comments**
**8. Figure 5: panel arrangement is confusing**
The arrangement of Figure 5 has been improved.

**9. Hoshii et al, 2024 is not listed in the references**
This omission has been corrected

**Reviewer #1 (Significance (Required)):**

**The key advance in this work is demonstration that Bod1L removal affects expression of DNA repair genes and increases DNA damage in ES cells. This seems to occur independently of changes in H3K4me3 (although specific analysis of H3K4me3 at these genes was not performed).**
We do now include analysis of H3K4me3 at promoters of genes downregulated after loss of Bod1L (new Supplemental Figure 6E). As with all active promoters, H3K4me3 is also elevated on these genes.
**Removal of Setd1a had similar effects. The work seems to support previous reports in leukemia cell lines for specific effects of Setd1a/Bod1L on DNA repair that is not related to Setd1a enzymatic activity (Hoshii et al 2018, 2024 in the manuscript)(and contrasts with findings in U2Os cells that do implicate enzymatic activity and emphasize a direct role at replication forks rather than in transcription; Higgs et al 2018 and Bayley et al 2022 in the manuscript). This is a modest but important conceptual advance in thinking about Bod1L and COMPASS complex functions in transcription and DNA damage repair. Whether these functions are specific to Bod1L vs Bod1 in ESCs, or to Setd1a over Setd1b, in ESCs, was not addressed.**
In the Introduction we highlighted the difference between Setd1a and b in ESCs. We previously published that Setd1a is essential whereas Setd1b is not, and Setd1b fails to rescue the loss of Setd1a in ESCs when (over)expressed from the Setd1a promoter (Bledau et al, 2014). Furthermore the preferential association of BOD1 with SETD1B rather than SETD1A can be concluded from the data provided. These considerations support the conclusion that SETD1A and BOD1L are specifically regulating DNA repair gene expression.
**The audience for this work will be chromatin/epigenetics experts. My expertise is in the field of chromatin and epigenetics, and I have studied histone modification function extensively in the context of transcription.**
As a final comment to reviewer 1 – thankyou for your thoughtfulness but please allow a comment on the term ‘COMPASS’, which is not used in our manuscript. COMPASS is a confusing and ambiguous term. It means ‘COMPlex ASsociated with yeast Set1’ and was first

used to describe the incomplete yeast Set1 complex. Concomitantly, my group published the complete complex, termed Set1C, along with the first biochemical proof that it was an H3K4 methyltransferase (the first bona fide H3K4 methyltransferase, and the second bona fide histone methyltransferase). The following year, the second COMPASS publication reported the full complex by including the missing subunit and also biochemical proof of H3K4 methyltransferase enzyme activity. Subsequently the term ‘COMPASS’ has been applied to the Trithorax and MLL complexes, which share half of the Set1C/COMPASS complex – these four subunits are highly conserved in eukaryotes – but also include another 4+ subunits unrelated to the other half of Set1C/COMPASS, which are also are different between the Trithorax/MLL1,2 and Trithorax related/MLL3,4 complexes. So it is imprecise and confusing – especially for the majority of bioscientists who do not have histone methylation expertise - to name these partially related but distinct complexes, ‘COMPASS’. The highly conserved 4 subunits of these complexes have been more precisely termed ‘WRAD’ (after the mammalian names, WDR5, RBBP5, ASH2L, DPY30). WRAD, as opposed to COMPASS, is unambiguous and does not require specialist insider knowledge to unravel the confusion. Therefore the term ‘WRAD’ is used to refer to the conserved quartet in the manuscript.

**Reviewer #2 (Evidence, reproducibility and clarity (Required)):**

**This manuscript investigates H3K4me-methylating complexes in mouse embryonic stem cells with a specific focus on the SETD1A complex and its binding partners. The authors define the interaction between SETD1A and BOD1L and characterize a requirement for BOD1L in maintaining the expression of DNA repair genes in mESCs. BOD1L has previously been implicated in regulating the replication fork (Bayley et al., Mol Cell, 2022 and others), however the authors propose another role for BOD1L in restraining H3K4me3 at TSS-proximal nucleosomes which impacts gene expression of DNA repair genes.**
With respect, we do not suggest that the restraining action of BOD1L on H3K4me3 has any impact on gene expression. In contrast, we note that elevated H3K4me3 at TSS-proximal nucleosomes does not correlate with changes in gene expression.

**However, a number of aspects of this model require additional support. Furthermore, while there are some new insights gained from the experiments performed, the data presentation makes the impact of the studies difficult to interpret. Specific concerns are outlined in further detail below:**
**1. A key approach used through the manuscript is AP-MS experiments to determine protein interactors of SETD1A, SETD1B, and other complex subunits. It appears these experiments were generally performed in triplicate, however, there should be more discussion of what thresholds were used to quantify interactors. The methods states that if 2 unique peptides were identified, though it is very difficult to tell from the volcano plots why some proteins are labelled and named as interactors and others are not discussed. Furthermore, the volcano plots are generally difficult to read and do not lend themselves well to comparisons between different experiments. Another format in addition to potential volcano plots, such as a**

**heatmap, would improve the readability and provide a better method of visualizing the quantitative results of these experiments.**
Our presentation of AP-MS data in Volcano plots is conventional. As mentioned above (reviewer 1, response 1), AP-MS analyses present candidate data that requires further support, which we supplied for the conclusions we draw, as detailed above.

**2. There is very little discussion of the additional interactors identified, outside of expected components, in the AP-MS experiments described in Figures 2 and 3. The authors state that they pursued additional experiments but that there was not productive data. It is unclear what this means as to whether these are not legitimate interactors or if there were other technical challenges. There is some discussion of OGT and BPTF, but it is not particularly informative. I think further clarification and/or characterization on the other interactors would be useful to be able to interpret the validity of the data presented in the AP-MS volcano plots.**
The candidate data obtained by AP-MS analyses include a core of reliable interactions as well as other less robust identifications that may be true or false positives. In this manuscript, we focused on the reliable and verified interactions, and mentioned notable additions, which we hope will assist further investigations. Further proteomic exploration is beyond the scope of this manuscript.

**3. In Figure 4, AP-MS is used to characterize interactors of different deletion mutants of SETD1A. The expression of the deletion mutants should be shown by western or another approach to see how these compare to wildtype. In addition, the interaction is further probed in cells by a co-IP approach using an overexpression construct of the X3 region of SETD1A. Since the authors have the deletion mutant, this could be used in a co-IP experiment in addition to exogenous expression of just the X3 fragment. This would allow a direct comparison with WT SETD1A and other mutants. Also, to further support the specificity of the interaction show in Fig 4D, a similar experiment could be performed with other regions of SETD1A (or B), such as a the X1 or X2 regions.**
We are not certain about these comments. The deletion mutants were examined by AP-MS, which is effectively superior to a co-IP, and retrieved most of the expected proteins. If we had pursued unexpected proteins identified in the mutant AP-MS, then Western (or similar) analysis of expression levels of the SETD1A mutants would be important. However we obtained reciprocal confirmation of the primary result, which is better than a co-IP with Western. Detailed analyses with other X regions are beyond the focus of this work.

**4. Figure 5C and this H3K4me3 chip results in the BOD1L mutant cells would be further supported by showing the levels of SETD1A (and other H3K4 methylating enzymes) in these cells lines to better support the conclusion that BOD1L is directly restricting SETD1A activity at chromatin.**
In both yeast and *in vitro*, elevated H3K4me3 by Set1C without Shg1 is not due to elevated Set1 expression or changes of the Set1 complex, (other than loss of Shg1; Roguev et al, 2001, Kim et al, 2013). Concordantly, we show that SETD1A without X3 (i.e. without BOD1L) still retrieves the

rest of the SETD1A-Complex (Figure 4B). Furthermore, Setd1a is expressed from its endogenous promoter to ensure physiological expression level.

**5. Figure 5F shows growth curves of WT and BOD1 mutant ESCs. However, this data requires statistical analysis to make an accurate comparison between cell lines. Furthermore, the mutant cell lines in particular would benefit from showing at least one additional time point if feasible. In addition, the authors state that this is likely representative of increased cell death in the mutant cells, however this is not directly tested in this experiment.**
Figure 5F shows straightforward growth curves of ESCs wt, heterozygous or homozygous Bod1l mutants. Please note that the figure includes the growth curves of two independent heterozygous and homozygous Bod1l ES cell lines thereby presenting reproducibility for the impaired growth.

**6. The volcano plots in Figure 6 for the RNA-seq analysis are also difficult to interpret. Another data presentation method should also be used to be able to compare between experiments- this is not that feasible with the method and labeling of the data here, and the quantitative aspect of this data is not fully realized using this approach.**
Figure 6A represents the RNA-seq data in Volcano plots, which is complemented by the dot plots of Figure 6B and the listing of genes in Table 1. RNA-seq data is difficult to present in visually accessible figures and we think that the presentations in Figure 6 are effective, because this visualization allows the estimation of effect size as well as p-values. These data are also presented in a different format in the Supplement Figure 6D.

**7. The authors propose that the role of BOD1L DNA damage repair in ESCs is two fold in ESCs-one is a direct role at replication forks, and a second is its role in regulation of DNA repair genes with SETD1A. This may be the case, but the data provided here do not show that there is a direct role for BOD1L in regulating these genes. Additional experiments showing chIP or CUT&RUN of BOD1L and or SETD1A-dependent H3K4methyl species would provide more evidence that these DNA repair gene expression changes are directly due to BOD1L's role. It is also possible the gene expression changes are an indirect consequence of it's role at replication forks, but this is not really addressed. Furthermore, the model proposed in Figure 8 is difficult to understand and does not clearly represent the data in places (for example, the impact on H3K4methylation).**
Our conclusion regarding the two-fold role of BOD1L is based on (a) the data of others regarding protection of the replication fork; (b) our verification that BOD1L is a component of the SETD1A complex; (c) the known role of SETD1A as the major H3K4 trimethyltransferase at active promoters; (d) the observation that conditional mutagenesis of Bod1l predominantly leads to decreased mRNAs that encode for various components of DNA repair pathways. If the loss of BOD1L led only to DNA damage and not gene expression changes due to compromised action of the SETD1A complex, then a loss of expression of DNA damage genes would not be expected. In particular, double strand DNA damage elevates the expression of Xrcc4, Xrcc5 and Rad51c however these mRNAs are strongly down-regulated when Bod1l (or Setd1a) is lost. These points constitute a strong basis for our conclusion of a two-fold role for BOD1L and these points have been strengthened in the revised manuscript.

Regarding the reviewers comments –
**This may be the case, but the data provided here do not show that there is a direct role for BOD1L in regulating these genes. Additional experiments showing chIP or CUT&RUN of BOD1L and or SETD1A-dependent H3K4methyl species would provide more evidence that these DNA repair gene expression changes are directly due to BOD1L's role.**

- it is extremely difficult to demonstrate a direct role for BOD1L in gene regulation using ChIP/CUT&RUN, because SETD1A and B, and subunits of their complexes, are found on all active promoters – (for example, Cxxc1; Fig. 3, Denissov et al 2014). The question of target gene specificity, which emerges from RNA-seq analyses, is a notable problem for the H3K4 methyltransferases because their widespread and overlapping chromatin occupancy on promoters does not facilitate conclusions about specificities. Stated differently, yes we find BOD1L on the affected promoters, but we also find BOD1L on all active promoters, so it's location on affected promoters is indecisive.

**8. The discussion section could be significantly streamlined and focused more directly on the content of the manuscript. There are a number of different areas covered in detail that go beyond what is needed for the discussion of the paper and are distracting and confusing. For example, the discussion of the work of Hoshii et al on page 13 is highly relevant, but it could be shortened to focus on the most relevant data from the 2024 paper. (Although I could not find this paper in the reference list, but assume it is this one: https://pubmed.ncbi.nlm.nih.gov/38989615/). Other areas that seems somewhat tangential include the discussion of the potential PP2A interaction on page 14, which is not focused on in the manuscript. Also, the broader question of the role of H3K4 methylation in transcription is covered in some detail and this could be shortened to discuss in a more straightforward manner the potential implications of this study on our understanding of H3K4methyl marks in transcription.**
With due respect, we disagree. A shorter, less informative and less thoughtful discussion would probably have been criticized as insufficient. The reviewer is expressing an opinion. We think that we have succinctly presented the complexities of H3K4 methylation in transcription and our brief comment about PP2A may assist further research.

**Reviewer #2 (Significance (Required)):**
**There are some new insights provided into the relationship between SETD1A/SETD1B and the BOD1 and BOD1L components of the complex, however, the overall advances of this manuscript are relatively limited. A number of additional experiments are required to advance this work beyond what is already known in the field, and the specificty of their conclusions needs additional support. This limits the overall impact of this study**
We thank the reviewer for acknowledging that there are some new insights. We have a different opinion regarding their impact on what is already known.

**Reviewer #3 (Evidence, reproducibility and clarity (Required)):**

**This manuscript reveals that BOD1L, as a subunit of the SETD1A complex, plays a key role in**

**embryonic stem cell survival by maintaining the expression of DNA repair genes to protect cells from the accumulation of DNA damage. The study finds that BOD1L interacts with the X3 helix of SETD1A through its Shg1 homology region, and that loss of BOD1L leads to elevated H3K4me2/3 levels (consistent with the conserved function of Shg1 in yeast), downregulation of DNA repair gene expression, accumulation of DNA damage, and cell death. While the findings possess a certain degree of novelty, there are some logical issues that require further revision.**

**1.The authors conclude that "BOD1L maintains DNA repair gene expression through the SETD1A complex," but the current evidence is merely correlational. However, does the downregulation of DNA repair genes directly lead to DNA damage accumulation and cell death? Does BOD1L's own function in replication fork protection (Higgs et al., 2015) also contribute to this phenotype? The authors mention this point in the discussion, but the experiments do not distinguish between these two functions.**

Please see the comments above responding to reviewer 2, point 7. The manuscript presents strong evidence that the conclusion is not 'merely coincidental'. We addressed the question '**Does BOD1L's own function in replication fork protection (Higgs et al., 2015) also contribute to this phenotype?** ' in the experiment of Figure 7 and not just mentioned in the discussion.

**It is recommended to supplement with rescue experiments: in BOD1L-deficient cells, complement with wild-type BOD1L and mutants (e.g., lacking the X3 binding domain) to examine DNA repair gene expression, DNA damage accumulation, and cell death.**

As noted in our response to reviewer 1 point 3, we thank the reviewer for this constructive suggestion for further experiments that ideally will be in another manuscript.

**2.Figure 5C shows that BOD1L deletion leads to increased H3K4me3, whereas SETD1A deletion results in decreased H3K4me3. However, RNA-seq reveals substantial overlap in the downregulated genes between the two conditions (Figure 6B). Based on this, the authors infer that "H3K4me3 is not essential for DNA repair gene expression." This inference is reasonable, but one possibility needs to be excluded: whether the increase in H3K4me3 caused by BOD1L deletion occurs at non-target genes (specifically, DNA damage repair genes). It is recommended to perform H3K4me3 ChIP-qPCR in BOD1L-deficient cells to validate changes at the promoter regions of key DNA repair genes.**

This comment is similar to a point made by reviewer 1 point 6. The analysis is now included in Supplement Figure 6E.

**3.Figure 5G uses γH2AX and pATM staining to detect DNA damage, but the type of DNA damage (double-strand breaks, single-strand breaks, replication fork stalling, etc.) and its extent have not been quantified. It is recommended to supplement with: (1) a neutral comet assay to detect double-strand breaks, or an alkaline comet assay to detect total DNA damage; (2) an analysis of replication fork stability (e.g., a DNA fiber assay) to distinguish between BOD1L's replication fork protection function and its transcriptional regulatory role.**

With respect, further DNA damage assays will not distinguish between BOD1L's replication fork protection function and its transcriptional regulatory role.

**4.Figure 2E shows that BOD1L interacts with BPTF independently of SETD1A. However, the functional significance of this interaction has not been further explored. BPTF is a subunit of the NURF chromatin remodeling complex, and its interaction with BOD1L may be involved in DNA repair or transcriptional regulation. It is recommended to supplement with: (1) examining changes in BPTF chromatin binding in BOD1L-deficient cells (BPTF ChIP-seq); (2) investigating whether BPTF knockdown affects DNA repair gene expression or the DNA damage response.**

The manuscript is not about BPTF, rather we present a complementary observation to assist further research.

**5.The study demonstrates that BOD1L binding to the X3 helix inhibits the methylation activity of the SET domain, but the molecular mechanism remains unclear. The authors propose hypotheses in the Discussion, such as "monomer vs dimer" or "allosteric regulation," but direct biophysical evidence to explain how this long-range regulation is achieved is lacking.**

The interaction between BOD1L and SETD1A is presented and the implications are discussed in the context of existing information on the Set1 complexes. Further work – indeed a completely new project - is required to explore the implications of the findings we report.

**6.The experiment observed that BOD1 can also bind to the X3 site of SETD1B, and the two proteins share structural similarity. Although the text mentions that BOD1L is the major subunit in ESCs, it does not sufficiently explore whether BOD1 exerts partial compensatory effects in the absence of BOD1L, or the logic underlying their specific switching in different tissues.**

We agree – the manuscript does not explore whether BOD1 exerts partial compensatory effects in the absence of BOD1L. That would also be another project. Also we have not included speculations about **'the logic underlying their specific switching in different tissues. '**

**Minor suggestion：**

**1.The RNA-seq experiments were performed with biological duplicates (two samples per condition), it is generally recommended to have at least three biological replicates to ensure statistical power.**

As specified in the M&M, the primary RNA-seq experiments were performed with biological duplicates in parallel in three closely related ESC culture conditions and the conclusions are drawn from these six overlapping datasets. The other RNA-seq experiment was performed in triplicates, as was an RNA-seq experiment using ESCFCS conditions, that was not included but delivered the same results as presented here.

**2.In the co-IP experiments shown in Figure 2D and 2E, there is a lack of quantification or internal controls.**

As mentioned in response to reviewer 1, point 1, the controls are included and quantification can be estimated from the figures. Further data are provided in Supplement Figure 1.

**3.The peak calling parameters and statistical methods for the ChIP-seq analysis were not described in detail.**

Now included in the M&M.

**4.The description of the BOD1L-BPTF interaction results (Figure 2E) in the main text is too brief, and the conditions and controls for the IP experiment are not specified.**

The text referring to BPTF has been expanded and is now –

Therefore, we examined the interaction with BPTF in more detail. By immunoprecipitation using BOD1L-VENUS expressed from a *Bod1l* BAC transgene, the interaction between BOD1L and BPTF was confirmed. However, immunoprecipitation using BPTF-VENUS expressed from a *Bptf* BAC transgene retrieved the NURF subunit SNF2L/SMARCA1 (Supplemental Fig. S1) but failed to retrieve SETD1A (Fig. 2E) indicating that BOD1L independently interacts with both SETD1A-C and BPTF, and that BPTF interacts with BOD1L independently of its interaction with NURF.

**5.The discussion section is somewhat lengthy and contains speculative content (such as the discussion on OGT and MLL complexes). Although interesting, these points are not strongly related to the core findings of this study and could be streamlined.**

The Discussion is a little less than 1300 words.

**6.Page8 "Bod1 esiRNAi knock-down" should be ""Bod1 esiRNA knock-down".**

Corrected.

**Reviewer #3 (Significance (Required)):**

**Should be revised.**

Revisons suggested by the reviewers have been incorporated.
